# A reversible SRC-relayed COX2-inflammatory program drives therapeutic resistance in BRAF^V600E^ colorectal tumors

**DOI:** 10.1101/2022.10.20.512885

**Authors:** Ana Ruiz-Saenz, Chloe E. Atreya, Changjun Wang, Bo Pan, Courtney A. Dreyer, Diede Brunen, Anirudh Prahallad, Denise P. Muñoz, Dana J. Ramms, Valeria Burghi, Danislav S. Spassov, Eleanor Fewings, Yeonjoo C. Hwang, Cynthia Cowdrey, Christina Moelders, Cecilia Schwarzer, Denise M. Wolf, Byron Hann, Scott R. VandenBerg, Kevan Shokat, Mark M. Moasser, René Bernards, J. Silvio Gutkind, Laura J. van ‘t Veer, Jean-Philippe Coppé

## Abstract

BRAF^V600E^ mutation confers a poor prognosis in metastatic colorectal cancer (CRC) despite combinatorial targeted therapies based on the latest understanding of signaling circuitry. To identify parallel resistance mechanisms induced by BRAF/MEK/EGFR co-targeting, we used a high throughput kinase activity mapping platform. We found that SRC kinases are systematically activated in BRAF^V600E^ CRC following targeted inhibition of BRAF ± EGFR, and that coordinated targeting of SRC with BRAF ± EGFR increases efficacy *in vitro* and *in vivo*. SRC drives resistance to BRAF ± anti-EGFR therapy independently of ERK signaling by inducing transcriptional reprogramming via beta-catenin (CTNNB1). The EGFR-independent compensatory activation of SRC kinases is mediated by an autocrine prostaglandin E_2_-loop that can be blocked with cyclooxygenase-2 (COX2) inhibitors. Co-targeting of COX2 with BRAF+EGFR promotes durable suppression of tumor growth in patient-derived tumor xenograft (PDX) models. COX2 inhibition represents a novel drug-repurposing strategy to overcome therapeutic resistance in BRAF^V600E^ CRC.

## INTRODUCTION

Presence of a BRAF^V600E^ kinase mutation predicts the worst prognosis form of metastatic colorectal cancer (mCRC). About 8% of mCRCs harbor a BRAF^V600E^ mutation. Because mCRC is the second leading cause of cancer death, it is estimated that more patients die of BRAF^V600E^ mCRC than melanoma each year. Unlike melanoma, BRAF^V600E^ mCRC does not respond to BRAF inhibitor monotherapy, and it responds only poorly to conventional chemotherapy ^1–3^. Response rates have increased by combining BRAF inhibitors with MEK inhibitors and/or an anti-EGFR antibody, however the majority of mCRC tumors still fail to regress and durability of disease control remains a challenge. Clinical outcomes have been remarkably similar with combinations of BRAF inhibitors (vemurafenib, dabrafenib, or encorafenib) and/or MEK inhibitors (trametinib or binimetinib) and/or anti-EGFR antibodies (cetuximab or panitumumab): with a median confirmed response rate of ~20%, progression free survival of ~4 months, and overall survival of ~9 months ^4–6^. In April 2020, the Federal Drug Administration (FDA) granted approval to the encorafenib plus cetuximab doublet for the treatment of patients with BRAF^V600E^ mCRC, based on comparable outcomes with this doublet vs. triplet combinations evaluated in clinical trials.

The objective of targeting BRAF ± MEK ± EGFR is to reinforce the inhibition of the main oncogenic driver pathway (BRAF–MEK) while jointly shutting down the activation of a drug resistance mechanism (EGFR) ^5,6^. However, the observed ceiling effect with this approach suggests that other prevalent mechanisms (i.e. dependencies) must cooperate to circumvent therapeutic effectiveness. The rational next step is to systematically identify orthogonal vulnerabilities, independent of the MAPK pathway, to inform the design of novel combination strategies to address this ongoing unmet medical need.

To test the hypothesis that other parallel pathways act as compensatory mechanisms to drug treatments, and to initiate an expanded search for complementary drug targets, we leveraged a high-throughput kinase-activity mapping (HT-KAM) platform. HT-KAM is a functional proteomic screening technology which enables direct measurement of the catalytic activity of many kinases in parallel ^7–10^. This systematic process can help identify the most significantly and specifically perturbed kinase hubs, in turn revealing actionable vulnerabilities (kinases or otherwise) that lie within phospho-circuits of cancer cells and tissues. Strategic kinase dependencies with the highest therapeutic potential can then be chosen for further investigation in cell culture and xenograft models. The ultimate goal is to identify rational therapeutic combinations capable of producing greater than incremental improvements in clinical outcomes for patients with BRAF^V600E^ mCRC.

We found that a highly conserved SRC-relayed inflammatory program drives the adaptive response to targeted therapies in BRAF^V600E^ CRC. Specifically, SRC family kinases were activated upon treatment with BRAF ± MEK ± EGFR inhibitors *in vitro* and *in vivo*, thus uncovering an EGFR-independent mechanism of resistance. We found that upon treatment with BRAF ± EGFR targeted therapy, the activation of SRC kinases regulates the downstream phosphorylation of beta-catenin (CTNNB1), which leads to the reprogramming of cells’ transcriptional profiles. Upstream of SRC, we found that SRC kinases were activated by an autocrine prostaglandin E_2_ (PGE2)-regulated GNAS-activation loop that COX2 inhibitors interrupted in both cell lines and patient derived xenograft (PDX) mouse models ^4,11^. This SRC-relayed mechanism of therapeutic resistance operated independently of ERK-signaling. We showed that supplementing the current standard-of-care combination of BRAF-inhibitor encorafenib plus an anti-EGFR antibody (panitumumab) with the FDA-approved COX2-inhibitor, celecoxib, significantly and consistently improved tumor growth inhibition. Overall, our study demonstrates that SRC signaling is at the nexus of a cell-autonomous inflammatory program with pro-tumorigenic activities, which explains why BRAF^V600E^ colorectal tumors develop resistance to BRAF/MEK and EGFR inhibitors. Our results suggest a new clinically actionable strategy, the addition of celecoxib to targeted therapies, to restore therapeutic response in BRAF^V600E^ CRC. This drug-repurposing approach is cost-effective with minimal added toxicity, and may be fast-tracked into clinical testing.

## RESULTS

### Kinome screening identifies SRC family kinases as independent functional determinants of the adaptive response to BRAF/MEK/EGFR targeting in BRAF^V600E^ CRC

We used the HT-KAM platform to measure kinase activity in extracts processed from WiDr cells, a well-established vemurafenib-resistant BRAF^V600E^ CRC model ^12–14^. Cells were treated with a BRAF inhibitor (vemurafenib) ± an EGFR inhibitor (gefitinib or cetuximab) for 8 hours. Peptide-level data (**Fig. 1a**) were then transformed into kinase activity signatures (**Fig. 1b**) using previously described deconvolution methods ^7^. We found that SRC displayed the most significantly increased kinase activity in response to BRAF inhibitor containing treatments (**Fig 1b**). Remarkably, the increase in SRC activity was conserved even following combined treatment with vemurafenib and either EGFR inhibitors (**Fig. 1b-d**), indicating that SRC is not a surrogate for EGFR-mediated resistance to BRAF targeted therapies.

**Fig. 1.**
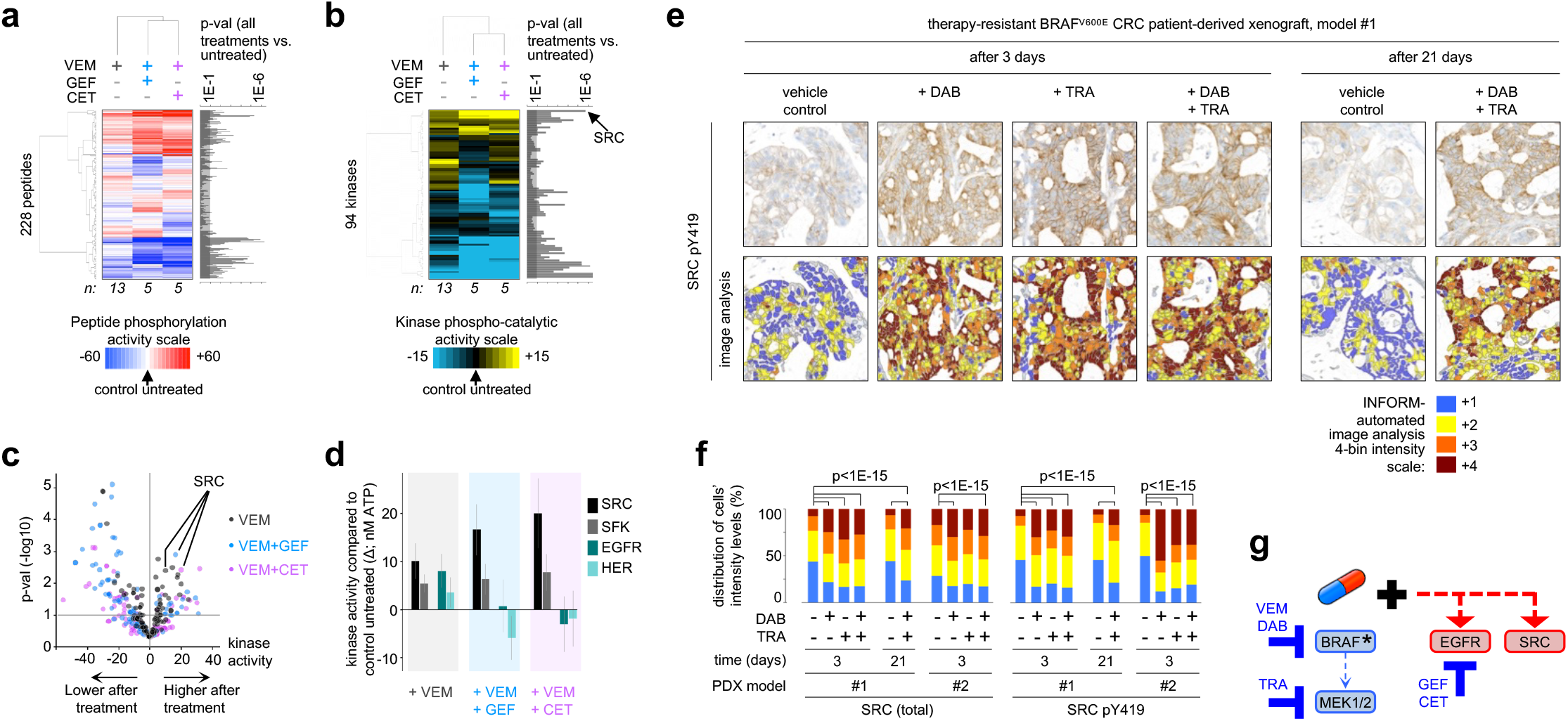
SRC is activated following BRAF/MEK/EGFR inhibition in BRAF^V600E^ CRC. **a**, **b**, Unsupervised hierarchical clustering of the phospho-catalytic activity signatures of WiDr cells treated with vemurafenib (VEM; n=13) ± gefitinib (GEF; n=5) or cetuximab (CET; n=5) as compared to their untreated control counterparts (n=23)**. a**, ATP consumption in cell extracts using 228 peptide sensors. **b**, Kinase signatures deconvoluted from peptide phosphorylation profiles in (**a).** Bar graphs next to the heatmaps display the p-values (p-val) for each of the peptides (**a**) or kinases (**b**) comparing all treated samples to controls. **c**, Volcano plot of data in (**b**) displays change in kinase activity versus p-values for each treatment arm. **d**, Bar graphs representing the shift in activity of SRC, SFK, EGFR and HER family kinases when cells are treated with VEM alone or in combination with GEF or CET. Kinase activity is compared to the untreated control cells and data displayed as the average ± standard error. **e**, Representative IHC images showing active SFK (phosphotyrosine-419 epitope in the SRC activation site) staining intensity following treatment of a BRAF^V600E^ CRC PDX model with vehicle control, dabrafenib (DAB) and/or trametinib (TRA) for 3 or 21 days. The color-coded bottom panel highlights differences in bin intensities resulting from automated image analysis. **f**, Quantification of IHC staining intensity for total and activated SFK in two PDX models treated for 3 or 21 days with DAB ± TRA vs. vehicle control. **g,** Proposed parallel mechanism of SRC activation in response to BRAF/MEK/EGFR therapies in BRAF^V600E^ CRC. BRAF*: BRAF ^V600E^.

SRC belongs to the SRC family kinase (SFK), which is composed of 11 membrane-associated, non-receptor tyrosine kinases that regulate cell proliferation, differentiation, apoptosis, migration, and metabolism among other processes ^15–18^. We validated the observed kinase activity signature by western blot in a panel of BRAF^V600E^ CRC lines after vemurafenib treatment. Results show increased SFK activation, as reported by the phosphorylation of Y419 (**Extended Data Fig. S1a**). We sought further substantiation of SRC-mediated resistance using rare xenograft models of BRAF^V600E^ CRC derived from patients who were subsequently treated with dabrafenib and trametinib on a clinical trial ^4^. PDX #1, and the corresponding patient’s tumor biopsy, exhibited primary resistance to treatment with dabrafenib + trametinib; whereas patient/PDX #2 exhibited early tumor regression and eventual progression. Using automated image analysis of IHC profiles, we observed statistically significant increases in active and total SRC in both PDX models after 3 or 21 days of treatment with dabrafenib and/or trametinib (**Fig. 1e,f** and **Extended Data Fig. S1b,c**). These data place SRC activation as an early adaptive response to BRAF inhibition, which is maintained even in residual tumors following BRAF and/or MEK inhibitor treatment. Unlike EGFR levels that were upregulated at baseline in BRAF^V600E^ cell lines ^12, 13^, comparison of untreated primary patient colorectal tumor specimens harboring or not a BRAF^V600E^ mutation showed that patient tumors start with similar levels of total SRC (**Extended Data Fig. S1d**). Together these findings led us to postulate that SRC is an EGFR-independent candidate drug target to overcome resistance to BRAF/MEK inhibitor therapies (**Fig. 1g**).

### SRC kinase inhibitors sensitize BRAF^V600E^ CRC cells to vemurafenib

To test the hypothesis that SRC is a druggable vulnerability in BRAF inhibitor resistant BRAF^V600E^ CRC, we first assessed the sensitivity of WiDr cells to two-drug combinations including a BRAF inhibitor and another kinase inhibitor, chosen based on the kinase signatures in **Fig. 1b**. Consistently, the greatest increase in cell growth inhibition and the highest combination index (CI) scores, i.e. synergy, were observed when the BRAF inhibitor was combined with a SRC inhibitor: dasatinib, saracatinib, or bosutinib (**Fig. 2a**; all CI>2.8). The potentiation in sensitivity to vemurafenib with the addition of a SRC inhibitor was conserved across various vemurafenib-resistant BRAF^V600E^ CRC cell lines (HT29, KM20, LIM2405, LS411N, OUMS23, RK01, SNUC5, VAC0432, WiDr), as shown in **Fig. 2b**; the EGFR-inhibitor gefitinib is included for comparison. This observation is specific to BRAF^V600E^ CRC, as it was not recapitulated in vemurafenib-sensitive BRAF^V600E^ melanoma cells (A375, A375 (SRC^Y530F^), A375 (myr-AKT1), Sk-Mel-28, Mel888) or BRAF wild-type CRC cells (HCT116, LoVo), as shown in **Fig. 2b** and **Extended Data Fig. S2a**. Furthermore, dual treatment with vemurafenib + dasatinib strongly inhibited colony formation in BRAF^V600E^ CRC cells, but not BRAF^V600E^ melanoma cells, at concentrations where dasatinib alone has a minimal effect (**Fig. 2c**). To further validate SRC’s role as a mediator of response to vemurafenib, we knocked down SRC using short hairpin RNA (shRNA) (**Fig. 2d**) and found that SRC-deficient BRAF^V600E^ CRC cells were more sensitive to vemurafenib treatment in 3-day viability and colony formation assays (**Fig. 2e-f**), although as expected not as profoundly as when using SFK inhibitors. Conversely, small interfering RNA (siRNA) knockdown of C-terminal SRC kinase (CSK), a negative regulator of SFK, led to increased SFK activation (**Extended Data Fig. S2b**) and reduced sensitivity to vemurafenib (**Extended Data Fig. S2c**). Collectively these data substantiate our hypothesis that SRC is a promising new target to overcome resistance to BRAF inhibitor therapies in BRAF^V600E^ CRC.

**Fig. 2.**
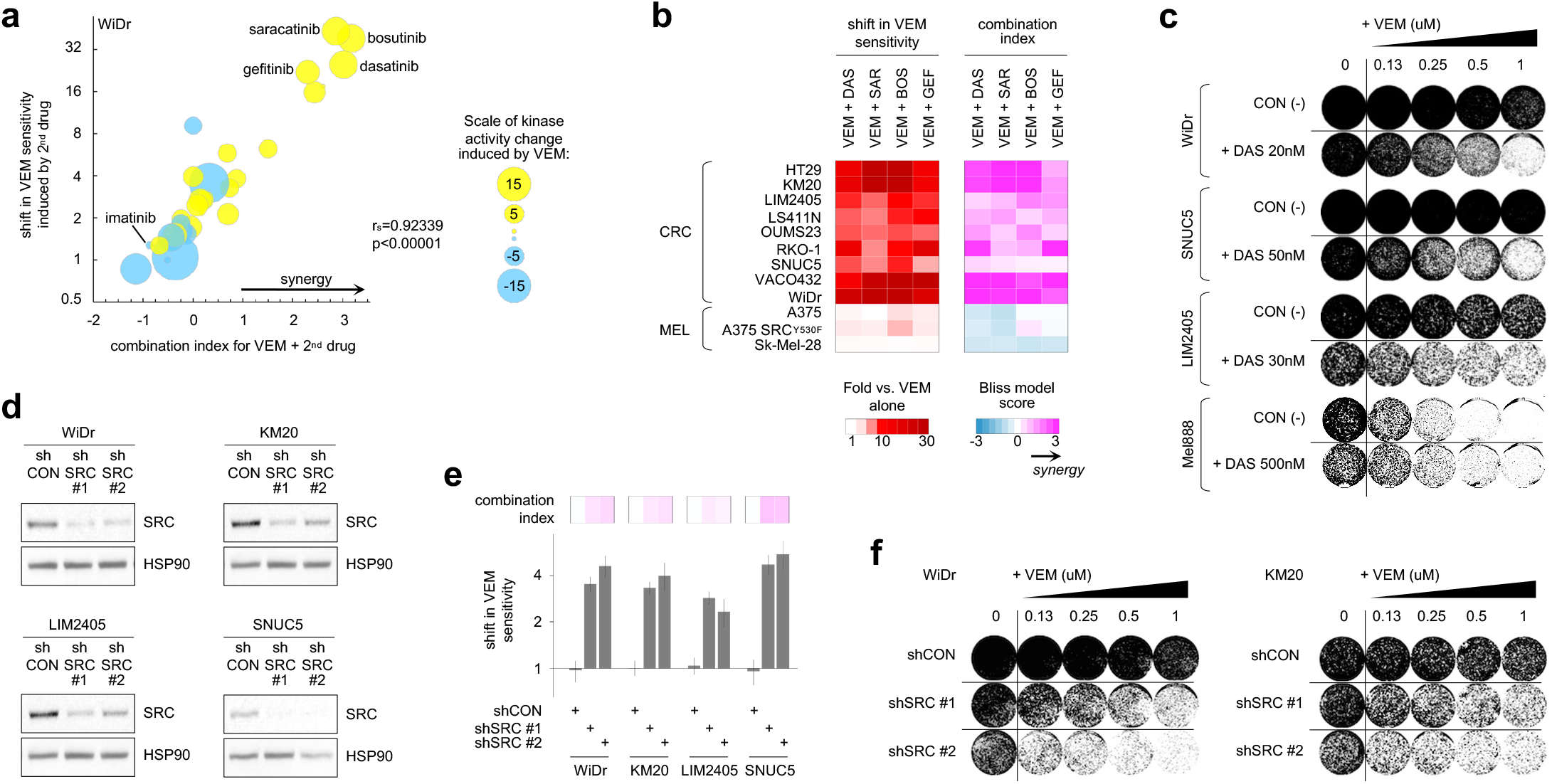
SRC kinase inhibition sensitizes BRAF^V600E^ CRC cell lines to vemurafenib. **a**, Cell viability assays evaluating WiDr cells treated with vemurafenib (VEM) plus a panel of kinase inhibitors selected based on results in **Fig. 1b**. The size of the bubble indicates the magnitude of the change in kinase activity induced by VEM treatment (**Fig. 1 b**), with color signifying increased (yellow) or decreased (blue) activity; y-axis: log2 scale; r_s_: Spearman’s rho correlation, p: p-value for 2-tailed t-test. **b**, Shift in VEM sensitivity, measured via cell viability assays (left panel) and calculation of the combination index (right panel) upon treatment of BRAF^V600E^ CRC or melanoma (MEL) cell lines with VEM together with a SRC inhibitor: dasatinib (DAS), saracatinib (SAR) or bosutinib (BOS), or an EGFR inhibitor: gefitinib (GEF), for three days. CI scores are averaged from individual experimental CIs calculated at 1x GI50, 2xGI50, 0.5xGI50 concentrations of each drug (n≥2). **c**, Colony formation assays where BRAF^V600E^ CRC or melanoma (Mel888) cells were treated with an increasing concentration of VEM alone (control, CON) or with a fixed dose of DAS. **d**, Western blot showing knockdown of SRC in BRAF^V600E^ CRC cell lines stably transfected with a control shRNA (shCON) or two different SRC-targeting shRNAs (shSRC). HSP90 serves as loading control. **e**, Bar graphs representing fold-change (log2 scale) ± standard error for change in sensitivity to VEM with knockdown of SRC in 3-day cell viability assays. Top panel: combination index, Bliss model score; colors as in **Fig. 1b. f**, Colony formation in BRAF^V600E^ CRC cells treated with an increasing concentration of VEM with or without SRC knockdown.

### SRC inhibition systematically improves efficacy of BRAF + EGFR targeting in BRAF^V600E^ CRC *in vitro* and *in vivo*

Since a BRAF + EGFR inhibitor doublet is the first FDA approved molecularly targeted regimen for patients with BRAF^V600E^ mCRC ^5^, we next asked whether the efficacy of BRAF and EGFR targeting can be improved by addition of a SRC inhibitor. To begin, we verified SFK activation after vemurafenib + gefitinib treatment in a panel of BRAF^V600E^ CRC cell lines (**Fig. 3a** and **Extended Data Fig. S3a**). The analysis showed increased SFK activation reflected by increased Y419 phosphorylation and Y530 de-phosphorylation, which further substantiates findings using HT-KAM in **Fig. 1b-d**. Moreover, triplet therapy with the addition of dasatinib to vemurafenib and gefitinib resulted in a synergistic increase in sensitivity to vemurafenib, greater than which was observed for either vemurafenib-containing doublet (**Fig. 3b** and **Extended Data Fig. S3b**). The synergistic impact on cell viability with the addition of a SRC inhibitor to BRAF + EGFR targeting was conserved across BRAF^V600E^ CRC cell lines, but not BRAF^V600E^ melanoma cells. Likewise, triplet treatment with vemurafenib plus gefitinib and dasatinib more effectively inhibited colony formation in BRAF^V600E^ CRC cells as compared to vemurafenib + gefitinib (**Fig. 3c**). Encouraged by these data, we tested combinations of BRAF ± EGFR and SRC inhibitors first in cell line-derived xenografts (**Fig. 3d**) and then in the same PDX models evaluated in Figure 1 (**Fig. 3e**). In all four xenograft models, triplet combinations resulted in statistically significantly improved tumor growth inhibition as compared to BRAF + EGFR or BRAF + SRC inhibitor doublets. Toxicity, as assessed by mouse weight and distress, was negligible (**Extended Data Fig. S3c,d**). Moreover, tumor regression was observed beyond the mid-point of treatment in multiple PDX tumors treated with dasatinib-containing regimens (**Extended Data Fig. S3e**). Next, we used a Generalized Linear Model (GLM) to quantify the effects of drug combinations vs. vehicle over time (**Fig. 3f-i**). Based on results in **Fig. 3d-i**, we found that the addition of a SRCi to a BRAFi had an equivalent or better effect on tumor growth inhibition in comparison to adding an EGFRi to a BRAFi. Furthermore, triple therapy with BRAFi + EGFRi + SRCi significantly improved tumor growth inhibition compared to any doublet (**Fig. 3f-g**; GLM standard coefficients >0.8 and >0.5 for cell line xenografts and PDXs respectively; FDR-corrected p-val in **Extended Data Fig. S3f** and in the right panel of **Fig. 3g**). **Fig. 3h-I** highlight that the addition of a SRCi to a BRAFi + EGFRi systematically and significantly improves effect size in both cell line and PDX models.

**Fig. 3.**
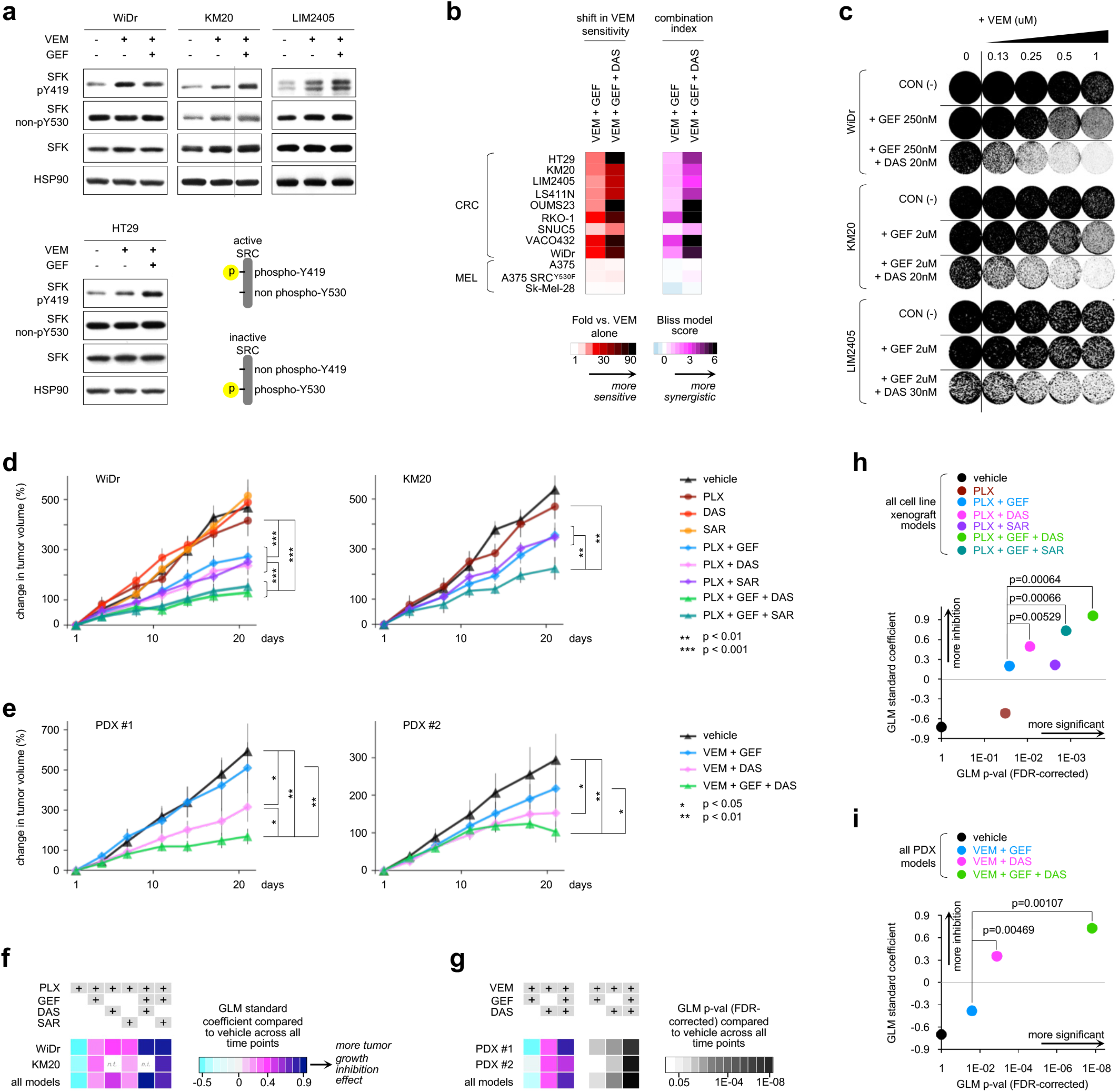
Coordinated targeting of SRC with BRAF + EGFR increases efficacy in BRAF^V600E^ CRC cell lines and xenografts. **a**, BRAF^V600E^ CRC cell lines treated with vemurafenib (VEM) ± gefitinib (GEF) were lysed and immunoblotted with the indicated antibodies. SFK activation is reflected by increased phosphorylation of the SRC activation site, tyrosine 419 (pY419), and non-phosphorylation of the inhibitory site, tyrosine 530 (non-pY530). Active SRC can be deactivated by re-phosphorylation of Y530 by C-terminal SRC kinase (CSK). HSP90 serves as loading control. **b,**Shift in VEM sensitivity measured via cell viability assays (left panel) and calculation of the combination index (right panel) upon treatment of BRAF^V600E^ CRC or melanoma (MEL) cell lines with VEM + GEF ± a SRC inhibitor, dasatinib (DAS), for three days. **c**, Colony formation assays where BRAF^V600E^ CRC cells were treated with an increasing concentration of VEM alone (control, CON) or with a fixed dose of GEF ± DAS. **d**, Treatment of cell line-derived xenograft mouse models with a VEM progenitor, PLX4720 (PLX); DAS; saracatinib (SAR); and/or GEF for 21 days. Plotted is the percent change in tumor volume relative to baseline (day 1). Data is displayed as the average for all mice in a specified treatment group ± standard error. **e**, Treatment of patient-derived xenograft models with VEM ± GEF ± DAS for 21 days, plotted as in **d**). **d-e**, All raw and relative tumor volumes can be found in the spreadsheets ‘Fig. 3d’ and ‘Fig. 3e’ of the **Source Data & Experimental Conditions**document. **f-g**, Generalized Linear Model (GLM) to test the association of change in tumor volume between treatment arms and vehicle over time. Effect size is measured as the GLM standard coefficient. GLM was applied to each tumor model separately or combined. Results for cell line xenografts and PDXs are respectively shown in **f** and **g**. GLM p-values corrected for false discovery rate (FDR) are shown in **g**. **h-i**, Comparison of effect size and FDR-corrected p-values of treatment arms with and without a SRC inhibitor.

### The activation of SRC upon BRAF ± EGFR inhibition regulates beta-catenin transcriptional reprogramming of BRAF^V600E^ CRC cells

Next, we sought to elucidate the mechanism underlying the synergistic effects of co-targeting SRC and the MAPK pathway ± EGFR shown in **Fig. 2**, **3**. MAPK signaling rebound is recognized as an important mechanism of resistance ^12, 13^, so we tested whether adding the SRCi dasatinib would inhibit phospho-ERK rebound more profoundly than would BRAFi alone or BRAFi + EGFRi. Using 8 BRAF^V600E^ CRC cell lines collected at 4 different times (up to 72h) with BRAFi or BRAFi + EGFRi or BRAFi + SRCi, we found that SRC inhibition did not significantly impact rebound of ERK phosphorylation (**Extended Data Fig. S4a, b**). This suggests that SRC activation in response to BRAF ± EGFR targeted therapy treatment acts via a distinct mechanism of drug resistance.

**Fig. 4.**
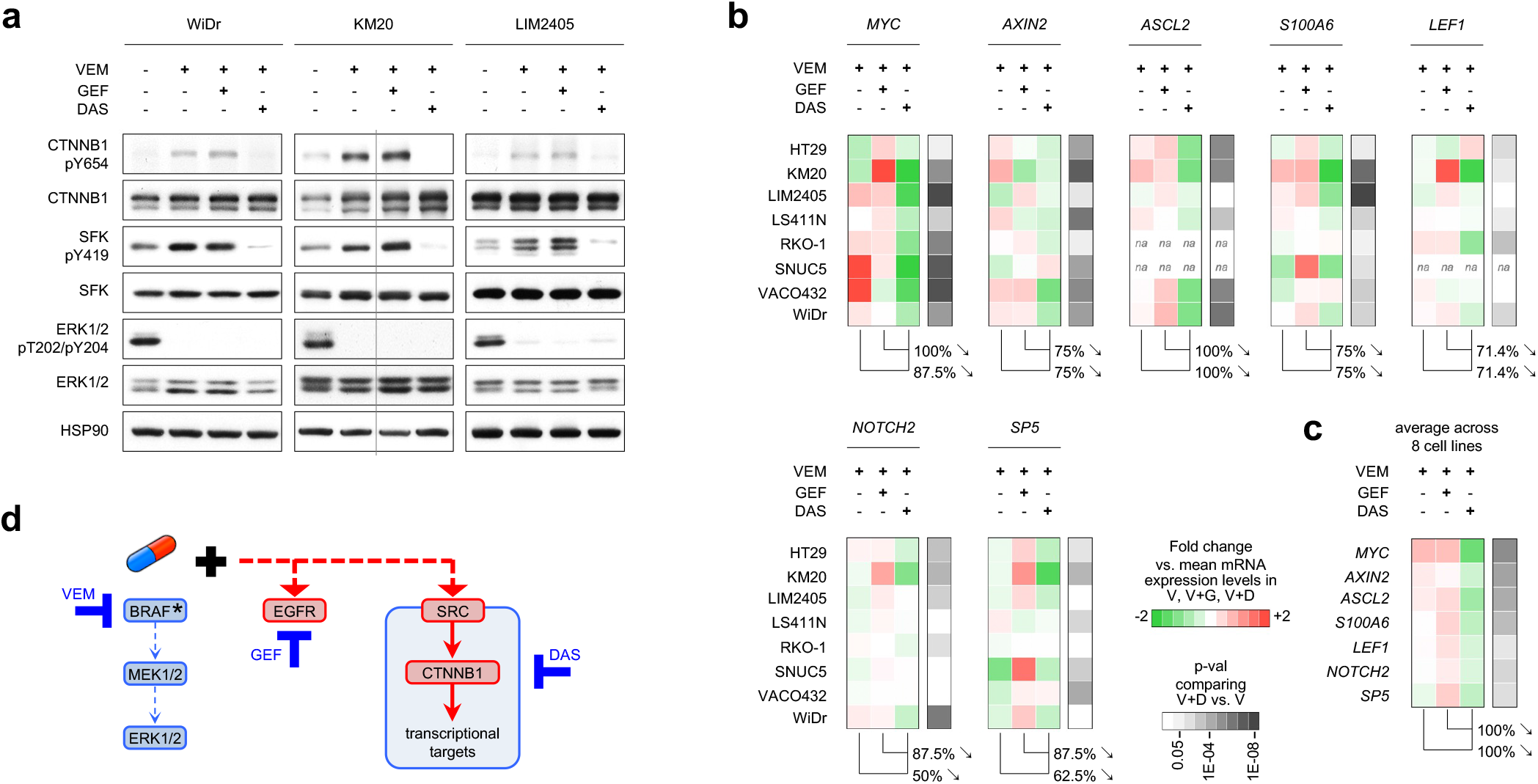
SRC regulates the phosphorylation of beta-catenin. **a**, Western blot of BRAF^V600E^ CRC cell lines treated with VEM ± GEF or DAS. The tyrosine 654 (Y654) of beta-catenin (CTNNB1) is a reported phospho-target site of SRC kinases ^8^. ERK1/2 phospho-T202/Y204 serves as a control for the effect of BRAF-inhibition (with VEM). SFK phospho-Y419 serves as a control for the effect of SFK-inhibition (with DAS). **b**, Color-coded expression levels of beta-catenin target genes (*MYC, AXIN2, ASCL2, S100A6, LEF1, NOTCH2, SP5*) measured using quantitative real time (qRT) PCR in BRAF^V600E^ CRC cell lines treated with VEM ± GEF or DAS. Expression profiles are shown as fold change against the mean mRNA expression level in VEM, VEM+GEF, VEM+DAS. Percentages indicate the proportion of measurements across 8 cell lines where the expression of the indicated gene (top) was lower with VEM+DAS than with VEM+GEF or VEM alone. The right-most columns indicate p-values (student t-test) comparing gene expression in VEM+DAS versus VEM alone. n.a.: not available due to expression levels that were too low. **c**, The expression profiles displayed in (**b**) were averaged across cell lines. **d**, Proposed mechanism regulated by SRC and that drives resistance to BRAF/MEK/EGFR therapies in BRAF^V600E^ CRC.

Since many kinases can propagate their pro-oncogenic activities via transcriptional re-programming, we hypothesized that SRC kinases may regulate the phosphorylation state of transcription factors involved in the compensatory response to BRAF ± EGFR inhibition. We first used the PhosphoAtlas kinase-substrate interaction database ^8^ to identify candidate transcription factor targets of SRC; we then assessed protein phosphorylation profiles by western blot. We found that the phosphorylation of beta-catenin (CTNNB1) at Y654, which is an understudied phospho-target site of SRC kinases ^56^, was increased upon BRAFi or BRAFi + EGFRi treatment, but was strongly decreased upon BRAFi + SRCi treatment (**Fig. 4a** and **Extended Data Fig. S4c**). To assess whether these changes in CTNNB1 Y654 phosphorylation impact the transcriptional activities of CTNNB1, we measured the expression levels of a series of CTNNB1 target genes ^57^ using quantitative RT-PCR. We found that adding the SRCi dasatinib to BRAFi led to a significant decrease in mRNA levels of all beta-catenin target genes we tested (i.e., *MYC, AXIN2, ASCL2, S100A6, LEF1, NOTCH2, SP5*) in comparison to BRAFi or BRAFi + EGFRi treated cells (**Fig. 4b,c**). This indicates that the activation of SRC kinases upon BRAF ± EGFR targeting, regulates the function of beta-catenin, a crucial transcription factor in CRC tumorigenesis, which can in turn re-program cells to sustain and adapt to therapeutic pressure. Altogether, SRC activation induces a tumor survival mechanism that acts in parallel to both the MAPK and EGFR signaling axes in BRAF^V600E^ CRC (**Fig. 4d**).

### In BRAF^V600E^ CRC, cyclooxygenase-2 (COX2) / prostaglandin E2 (PGE2) upregulation drives BRAF inhibitor-induced SRC activation *in vitro* and *in vivo*

Acknowledging that addition of a SRC inhibitor to BRAF ± MEK ± EGFR targeted therapies for treatment of BRAF^V600E^ mCRC is unlikely to be clinically acceptable due to concerns about toxicity in patients, we asked whether characterization of upstream effectors of SFK could lead to a more clinically appropriate regimen. It has been reported previously that prostaglandin E2 (PGE2) can induce activation of SRC in CRC cells, without specific attention to BRAF^V600E^ mutation status ^19, 20^. Using a panel of BRAF^V600E^ CRC cell lines, an increase in PGE2 levels in the media was consistently found as a result of BRAF ± EGFR inhibition (**Fig. 5a**). Treatment of BRAF^V600E^ CRC cells with PGE2 led to an increase in SFK activation (**Fig. 5b**), and a 2 to 4-fold increase in resistance to vemurafenib, with CI scores indicating antagonism (**Fig. 5c**). Consistent with the SRC / beta-Catenin signaling cascade established in **Fig. 4**, we found that PGE2 treatment led to an increase in phosphorylation of CTNNB1 Y654 (**Extended Data Fig. S5a**).

**Fig. 5.**
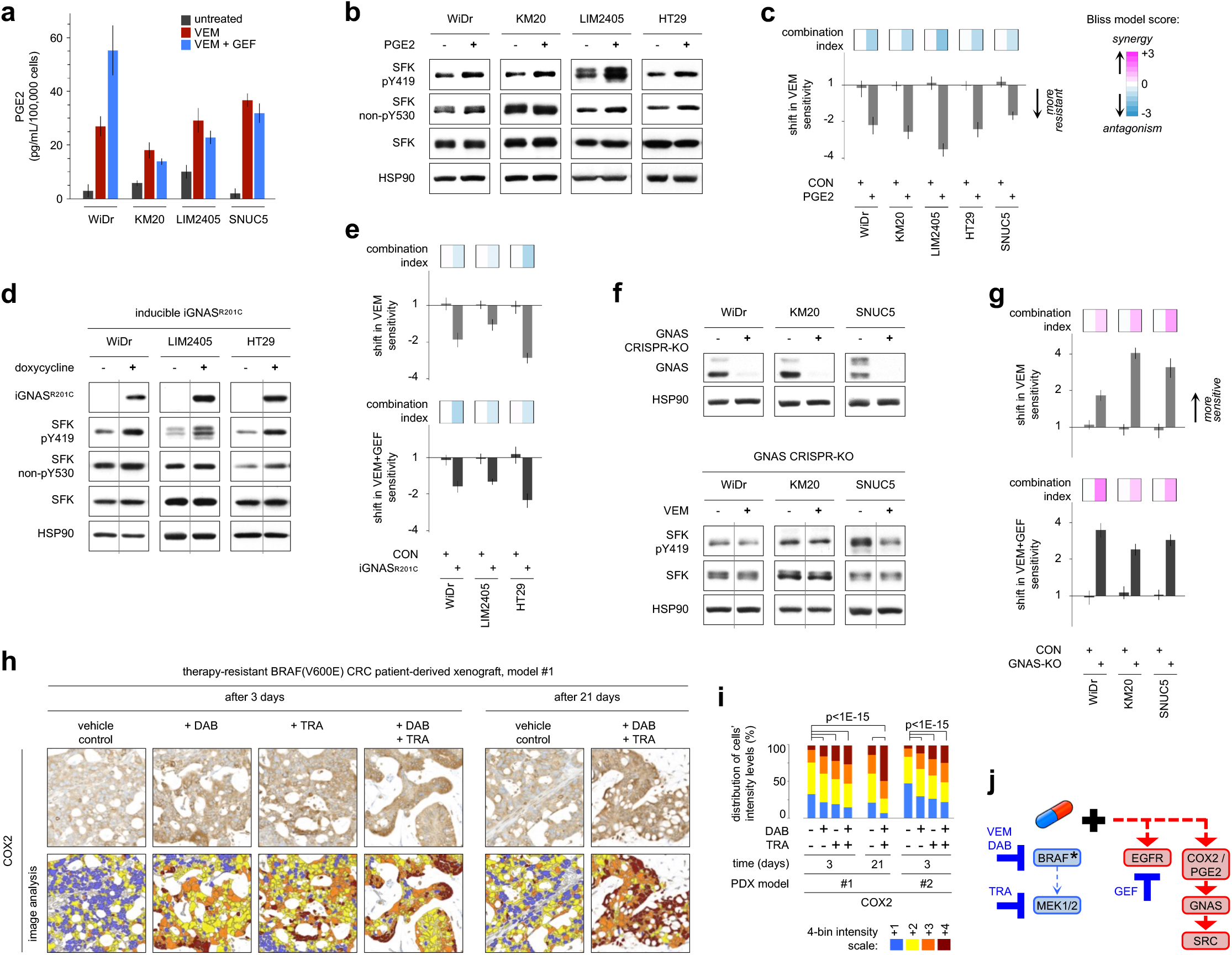
COX2/PGE2 upregulation mediates SRC activation in BRAF^V600E^ CRC cell lines and PDXs. **a**, PGE2 secreted levels were measured by ELISA in the conditioned media of BRAF^V600E^ CRC cell lines treated with vemurafenib (VEM) ± gefitinib (GEF). Data is displayed as the average PGE2 secretion in pg/mL per 100,000 cells ± standard deviation**. b**, BRAF^V600E^ CRC cell lines were treated with exogenous PGE2. Cell lysates were assayed by western blot as indicated. Y419 phosphorylation and lack of phosphorylation of Y530 (non-pY530) are used as readouts of SFK activation. HSP90 serves as loading control. **c,** Bar graphs representing fold-change (log2 scale) ± standard error for change in sensitivity to VEM upon further treatment with PGE2 or untreated control (CON) in 3-day cell viability assays. Top panel: combination index, Bliss model score. **d**, Three BRAF^V600E^ CRC cell lines engineered with a doxycycline-inducible constitutively active GNAS construct, iGNAS^R201C^, were treated with doxycycline. Cell lysates were assayed by western blot as indicated. **e**, Bar graphs representing fold-change (log2 scale) ± standard error for change in sensitivity to VEM or VEM+GEF after iGNAS^R201C^ induction in 3-day cell viability assays. Top panel: combination index, as in **Fig. 5c. f**, GNAS was knocked out in BRAF^V600E^ CRC cells using CRISPR (GNAS-KO). GNAS-KO was validated by western blot (top panel). GNAS-KO cells were treated with VEM and cell lysates were assayed by western blot with the indicated antibodies (bottom panel). **g,** Bar graphs representing fold-change (log2 scale) ± standard error for change in sensitivity to VEM or VEM+GEF with GNAS-KO in 3-day cell viability assays. Top panel: combination index, as in **Fig. 5c. h**, Representative immunohistochemistry (IHC) images showing COX2 staining intensity following treatment of a BRAF^V600E^ CRC PDX model with vehicle control, dabrafenib (DAB) and/or trametinib (TRA) for 3 or 21 days. The color-coded bottom panel highlights differences in bin intensities resulting from automated image analysis. **i**, Quantification of COX2 staining intensity by IHC for two PDX models treated for 3 or 21 days with DAB ± TRA vs. vehicle control. **j**, Proposed mechanism of COX2/PEG2 mediated SRC-driven resistance to BRAF/MEK/EGFR therapies in BRAF^V600E^ CRC.

High levels of PGE2 promote tumor growth by eliciting aberrant extracellular signaling via its G-protein-coupled receptors (GPCRs), EP2 and EP4, and their key downstream effector, GNAS ^19, 21–23^. To mimic the effect of PGE2, we engineered three BRAF^V600E^ CRC cell lines with doxycycline-inducible expression of a constitutively active GNAS mutant (GNAS^R201C^). When these cells were treated with doxycycline, GNAS^R201C^ was induced, leading to increased SFK activation (**Fig. 5d**). Induction of GNAS also rendered the cells more resistant to vemurafenib ± gefitinib treatment; CI scores again indicate antagonism (**Fig. 5e**). On the other hand, suppressing GNAS expression by CRISPR-knock out GNAS in BRAF^V600E^ CRC cell lines (GNAS-KO) prevented SFK activation after vemurafenib treatment (**Fig. 5f**). GNAS-KO cells displayed increased sensitivity to BRAF ± EGFR inhibition, with CI scores showing synergy (**Fig. 5g**), indicating that treatment-dependent SFK activation in BRAF^V600E^ CRC cells is downstream of PGE2/GNAS signaling. We noted that neither GNAS^R201C^-induced activation of SFK nor GNAS-CRISPR knockout-induced inhibition of SFK activity impacted the rebound of ERK phosphorylation (**Extended Data Fig. S5b-c**), further indicating that the feedback activation of the PGE2–GNAS–SRC signaling axis acts in concert with the MAPK-cascade to drive therapeutic resistance.

It is well established that PGE2 expression and secretion are regulated by COX2 ^58^. COX2 upregulation in BRAF^V600E^ CRC PDX tumors treated with dabrafenib + trametinib for 3 or 21 days was corroborated by IHC (**Fig. 5h**, **i**). Paralleling what was observed with SRC in these same PDX persists in residual tumors even at late time points. In untreated primary patient colorectal tumor specimens with or without a BRAF^V600E^ mutation, COX2 levels were similar (**Extended Data Fig. S5d**), again corresponding to observations with SRC (**Extended Data Fig. S1d**). In summary, these findings suggest that, in BRAF^V600E^ CRC, the COX2/SRC/beta-catenin signaling does not overlap with the EGFR/BRAF/MAPK signaling, and it is the treatment with BRAF ± MEK or EGFR targeted therapies that triggers the compensatory upregulation of a pre-existing COX2-PGE2-GPCR-GNAS autocrine loop, which in turn activates SRC (**Fig. 5j**).

### COX2 inhibition synergistically improves efficacy of BRAF/MEK/EGFR targeting in BRAF^V600E^ CRC *in vitro* and *in vivo*

COX2 is a rational drug target, given its robust association with CRC tumor progression in patients ^24^, although no prior clinical trials have focused on BRAF^V600E^ CRC. COX2 inhibitors also represent a practical alternative to SRC-targeting therapies: the COX2 inhibitor, celecoxib, is FDA-approved, has a favorable side-effect profile and is relatively inexpensive. Thus, the logical next step was to test BRAF-targeted therapies in combination with COX2 inhibition. As proof of concept, addition of a COX2 inhibitor produced a consistent, synergistic increase in sensitivity to vemurafenib across a panel of BRAF^V600E^ CRC cell lines, which was not recapitulated in BRAF^V600E^ melanoma (**Fig. 6a**). Using very low individual drug concentrations of trametinib, gefitinib and celecoxib (GI10 or below), we were able to demonstrate potentiation of cell growth inhibition with combinations of up to four targeted therapies (**Fig. 6b**, left panel). The addition of celecoxib systematically improved the efficacy of –and synergized with– vemurafenib + trametinib ± gefitinib (**Fig. 6b**, right panel). The most synergistic combination, across BRAF^V600E^ CRC but not melanoma cell lines, was the quadruple treatment arm.

**Fig. 6.**
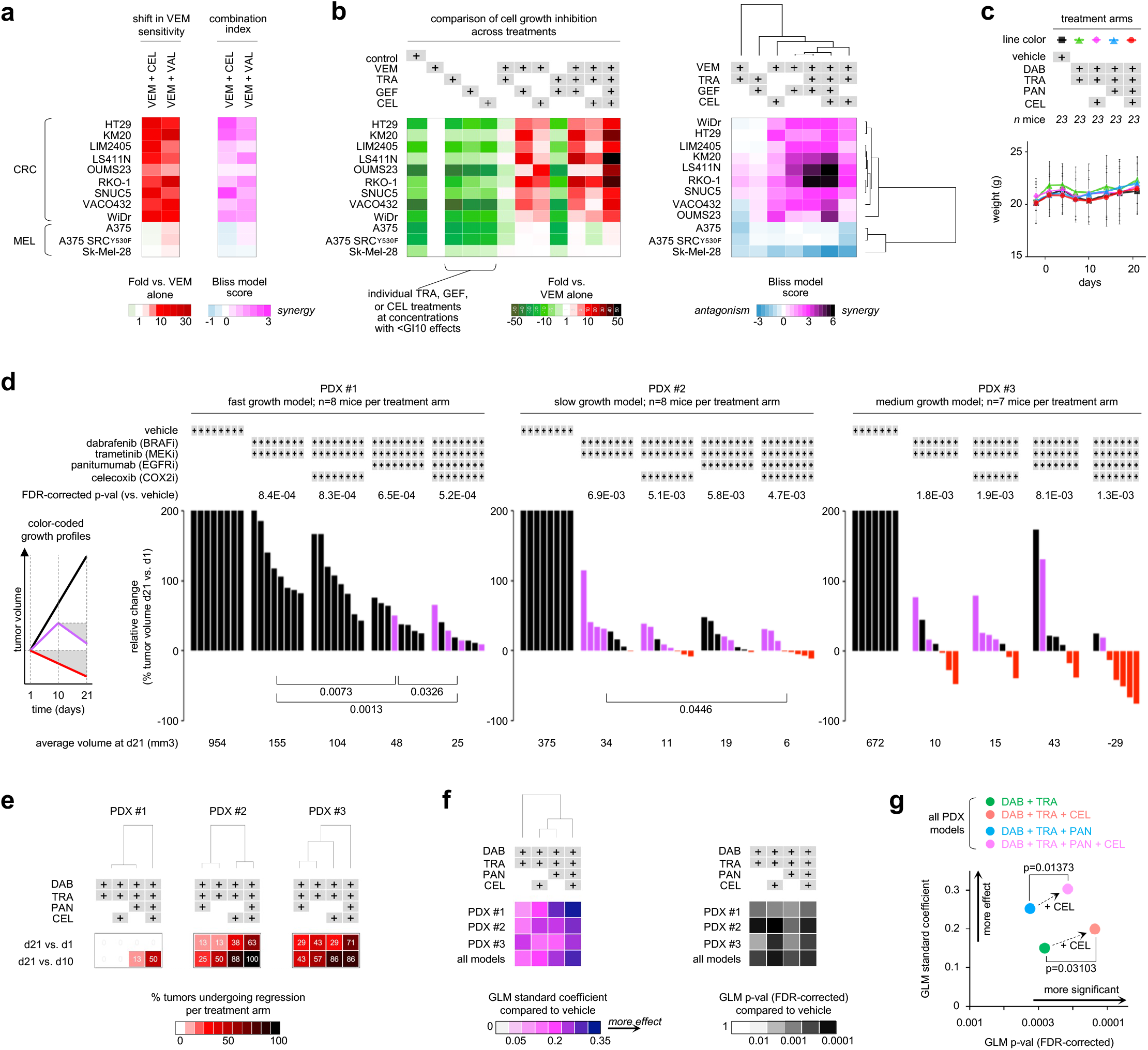
Coordinated targeting of COX2 with BRAF/MEK/EGFR improves efficacy in BRAF^V600E^CRC cell lines and PDXs. **a**, Shift in vemurafenib (VEM) sensitivity measured via cell viability assays (left panel) and calculation of the combination index (right panel) upon treatment of BRAF^V600E^ CRC or melanoma (MEL) cell lines with VEM together with a COX2 inhibitor: celecoxib (CEL) or valdecoxib (VAL), for three days. CI averaged from experimentally measured CIs at 1×GI50, 2×GI50, 0.5xGI50 concentrations of each drug (n≥2). **b**, Treatment of BRAF^V600E^ CRC or MEL cell lines with up to four inhibitors: TRA, GEF and CEL at concentrations that result in at most 10% of maximal inhibition of cell proliferation (GI10) when dosed individually. Cell growth inhibition across treatments permutations, normalized to VEM monotherapy (left panel), was used to calculate the combination index relative to all other treatment arms and subjected to unsupervised hierarchical clustering comparing cell lines and treatment arms (right panel). **c**, Mouse weight as a surrogate for toxicity following treatment of BRAF^V600E^ CRC PDXs (23 mice per treatment arm) with vehicle control, dabrafenib (DAB), trametinib (TRA), celecoxib (CEL) and/or PAN (panitumumab). Data is displayed as the average weight in grams ± standard deviation. **d**, Tumor growth inhibition in BRAF^V600E^ CRC PDX models following treatment with DAB +TRA ± CEL ± PAN or vehicle (control). Waterfall plots show the relative change in tumor volume: each bar represents one tumor; and the height of the bar compares the final volume at day 21(d21) to the starting volume at day 1. Volume changes are capped at 2-fold of the starting volume (i.e. 200%). Tumors that regressed by day 21 are shown in red (compared to volume at day 1) and purple (compared to volume at mid-treatment, i.e. day 10). Average final tumor volumes per treatment group are indicated underneath the graph (black font). T-test p-values are indicated when p<0.05. All raw and relative tumor volumes can be found in the spreadsheet ‘Fig. 6d’ of the **Source Data & Experimental Conditions**document. **e**, Semi-supervised hierarchical clustering of the percentages of regressing tumors per treatment arm are indicated, comparing day 21 vs. day 1 and day 21 vs. day 10. **f**, Generalized Linear Model (GLM) to test the association of change in tumor volume between treatment arms and vehicle over time. GLM was applied to each PDX model and all PDX combined. Left panel, effect size measured as the GLM standard coefficient; semi-unsupervised hierarchical clustering further compares the efficacy of the treatment arms. Right panel, ranking by GLM p-values corrected for false discovery rate (FDR). **g**, Comparison of effect size and FDR-corrected p-values of treatment arms with and without the addition of celecoxib.

Next, we tested whether addition of celecoxib could improve upon two clinical benchmark regimens: the dabrafenib + trametinib doublet received by the patients from whom the PDX models were derived ^4^, and a triplet regimen with addition of the anti-EGFR antibody, panitumumab, which was tested in a subsequent clinical trial ^6^. Toxicology studies were conducted prior to efficacy testing: mouse weight, a surrogate for drug toxicity, remained stable over the course of therapy for all inhibitor combinations (**Fig. 6c**). The critical finding from the efficacy studies was that addition of celecoxib to the clinical trial-tested doublet and triplet drug regimens resulted in consistently superior tumor growth inhibition in all three BRAF^V600E^ CRC PDX models (**Fig. 6d**). The majority of tumors treated with celecoxib in addition to dabrafenib + trametinib ± panitumumab exhibited regression by the second half of the 21-day treatment course (**Fig. 6d-e**). It is notable that 78% of tumors subjected to quadruple therapy with celecoxib exhibited regression (red and purple bars), vs. 30% of tumors regressing in the no-celecoxib triplet therapy counterpart arm. Across models, quadruple treatments resulted in the most statistically significant tumor growth inhibition (**Fig. 6d**; FDR-correct p-values above the bar graphs).

When applying the GLM approach across PDX models, we found that the addition of celecoxib systematically increased effect size; the greatest effect size was observed for the quadruple therapy (**Fig. 6f**, left panel; **Fig. 6g**, y-axis). Moreover, addition of celecoxib to any treatment arm resulted in statistically significant improvements in tumor growth inhibition (**Fig. 6f**, right panel; **Fig. 6g**, x-axis and arrows).

### Addition of celecoxib to current standard of care treatment results in durable tumor growth inhibition in BRAF^V600E^ CRC PDXs

Findings in **Fig. 6d** prompted us to assess the durability of treatment effects in the two most drug resistant PDXs (PDX #1 and #2 in **Fig. 6d**). We measured changes in tumor volume for >50 days in mice treated with encorafenib (BRAFi) ± panitumumab (EGFRi) ± celecoxib (COX2i) (**Fig. 7a-b**). We found that the triple treatment significantly improved tumor growth inhibition as compared to the dual drug combination, which is a current standard of care ^5^ (p-values across measurements underneath the graph in **Fig. 7a-b**). This was confirmed by GLM analysis considering all time points and individual tumor volumes (**Fig. 7c-d**). Increased toxicity was not observed with the addition of celecoxib (**Fig. 7e**). These results indicate that COX2 inhibition represents a novel, low cost and low toxicity drug-repurposing strategy to overcome therapeutic resistance in BRAF^V600E^ CRC: supplementing encorafenib + panitumumab with celecoxib durably improves tumor growth inhibition.

**Fig. 7.**
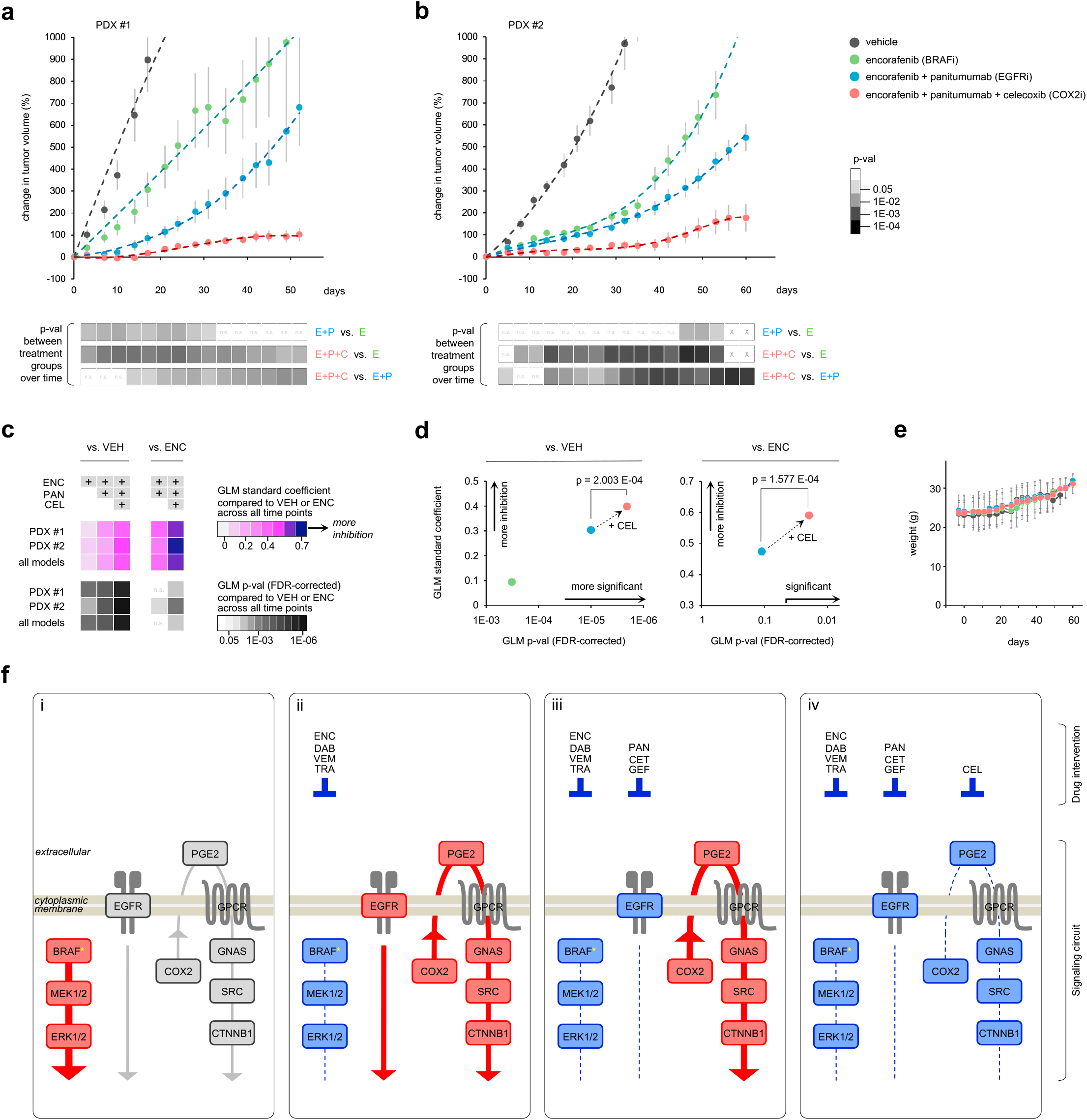
Coordinated targeting of COX2 with BRAF + EGFR improves long term efficacy in BRAF^V600E^ CRC PDXs. **a-b**, Tumor growth profiles in BRAF^V600E^ CRC PDX models #1 and #2 treated with ENC ± PAN ± CEL or vehicle (control). Changes in tumor volumes relative to starting volume at day 1 (average ± standard error) are plotted over time. Student t-test p-values across all time points comparing treatment arms are shown as a grey scale underneath each graph. n.s. : not significant; x : not available. All raw and relative tumor volumes can be found in the spreadsheets ‘Fig. 7a’ and ‘Fig. 7b’ of the **Source Data & Experimental Conditions** document. **c**, GLM to test the association of change in tumor volume over time, either between treatment arms and vehicle (left panel) or between combination therapy and ENC alone (right panel). GLM was applied to each individual PDX model and to both PDXs combined. The top section shows effect size measured as the GLM standard coefficient comparing the efficacy of the treatment arms. The bottom section shows GLM FDR-corrected p-values. **d**, Comparison of effect size and FDR-corrected p-values of treatment arms with and without celecoxib, using vehicle or ENC treatment as the baseline (respectively left and right). **e**, Mouse weight as a surrogate for treatment toxicity. Data is displayed as the average weight in grams ± standard deviation. **f**, Schematic summary of the states of signaling pathways depending on treatment: (i) untreated tumors, with BRAF-MEK-ERK as the main driver of progression (red), and baseline activity of the EGFR and COX2 / SRC signaling pathways (grey); (ii-iv) tumors treated with drugs (listed on top) inhibiting the activity (blue) of the three distinct signaling axes: BRAF-MEK, EGFR and COX2-SRC-beta Catenin; (iv) triple treatment collectively blocks the cooperative dependencies that drive resistance and progression.

## DISCUSSION

Despite recent optimization of targeted therapy combinations ^3–6^, BRAF^V600E^ still predicts the worst prognosis form of mCRC. Thus, we endeavored to uncover orthogonal mediators of compensatory resistance to BRAF, MEK and EGFR targeted inhibitors tested in patients. We discovered that SRC kinases act as a nexus of adaptive, druggable, and EGFR-independent therapeutic resistance *in vitro* and *in vivo*. Our findings were reproducible across a variety of inhibitors in the same class, cell lines, and mouse models, and yet were specific to BRAF^V600E^ mCRC. The activation of SRC in response to BRAF ± MEK or EGFR therapies did not contribute to MAPK signaling rebound. Instead, we found that SRC activation regulates transcriptional reprogramming via beta-Catenin activation, and is mediated by an upstream pro-inflammatory pathway involving COX2. Addition of celecoxib to inhibitor combinations tested in clinical trials consistently resulted in superior and durable tumor growth suppression in BRAF^V600E^ CRC PDX models. Our study identified unanticipated cooperative dependencies of actionable targets, yielding new strategies to overcome therapeutic resistance (summarized in **Fig. 7f**).

BRAF^V600E^ is a classic example of how the same activating mutation can play different roles depending on the cancer subtype-specific signaling context. BRAF inhibitors have produced an impressive response rate in BRAF^V600E^ melanoma— but not in CRC ^3^. In CRC, synthetic lethality genetic dropout screens originally found that feedback activation of EGFR promotes intrinsic resistance to BRAF inhibition ^12–14, 25^. However, modest responses in patients treated with BRAF + EGFR combination therapy underline why, in situations where predicting therapeutic response cannot be reduced to a single genetic dependency, there is a role for functional proteomic approaches designed to more comprehensively reveal crosstalk between signaling pathways, and to better dissect the dynamic processes induced by drug interventions ^26–29^. Here, we used the HT-KAM platform to directly capture the phospho-catalytic fingerprint of kinases in biological extracts, and to identify ranked drug susceptibilities. This elucidated how the concerted rewiring of interdependent signaling pathways drives resistance to BRAF, MEK and EGFR targeted therapies. SRC was identified as a central, conserved mediator of these signaling circuits in BRAF^V600E^ CRC.

SRC kinases are a non-receptor protein tyrosine kinase (NRTK) family of essential pleiotropic mediators of signaling cascades that connect extracellular cues to intracellular programs ^17, 18, 30^. Knowing that SRC often acts downstream receptor protein tyrosine kinases (RTKs), including in the context of acquired resistance to RAF inhibition ^59–60^, one might expect SRC to be activated by EGFR in response to BRAF-targeted therapy ^12,13,25^. Unexpectedly however, in BRAF^V600E^ CRC, we found that SRC kinases function as integral components of a drug-resistance circuit that is triggered independently of EGFR. Specifically, BRAF^V600E^ CRC cells adapt to targeted therapy by relying on a separate inflammatory loop that funnels through SRC. Possibly even more surprising, our *in vivo* and *in vitro* observations (**Fig. 3**) indicate that SRC may have a more dominant role than EGFR in the context of BRAF^V600E^ CRC. In fact, previously discovered transactivation effects of SRC onto EGFR via intra- and extra-cellular mechanisms ^21,31^ show that SRC acts as an upstream effector of EGFR. A SRC-driven transactivation mechanism would provide an alternative route to the previously noted feedback-release activation of EGFR via reduced CDC25C phosphatase activity that is initiated by therapeutic inhibition of BRAF/MEK ^13^. Together, CDC25C and SRC could functionally complement each other by converging on EGFR to coordinate its activation upon BRAF/MEK-inhibition.

Although SRC kinases are known direct upstream effectors of c-RAF, which regulate the signaling activity of RAF homo-/hetero-dimers ^32–35^, inhibition of SRC did not impact the ERK rebound commonly associated with RAF-therapy resistance. A SRC-driven drug-bypass mechanism might still explain how tumors can efficiently evade RAF-targeting, without acquired resistance mutations. Moreover, SRC kinases can directly promote the activity of other kinases involved in drug resistance, including AKT1 ^36^, an essential mediator of EGFR-signaling. BRAF^V600E^ CRC cells may thus compensate for the strain of EGFR-targeting via SRC, releasing cells from their dependency on EGFR to adapt to BRAF + EGFR combination therapies.

In addition to these direct regulatory effects on downstream kinases, SRC kinases are also known to propagate their pro-oncogenic activities via networks of transcription factors ^37–39^. In fact, we found that SRC phosphorylates an understudied phospho-site of beta-Catenin (Y654). Upon BRAF ± EGFR inhibition, SRC-dependent phosphorylation of Y654 increases beta-Catenin’s transcriptional activity, leading to a reprogramming of the transcriptional profiles of BRAF^V600E^ CRC cells. The WNT/beta-catenin signaling pathway is a key driver in the initiation and progression of CRC, differentiating it from other cancers including BRAF-mutated melanoma. Such SRC-driven reprogramming mechanism could rapidly and durably re-wire signaling pathways in BRAF^V600E^ CRC cells, effectively promoting adjustment to therapeutic stress.

The signaling plasticity offered by these SRC-dependent mechanisms may usurp and/or reinforce cell-autonomous pathways that drive tumor survival while bypassing other dependencies, including under the influence of BRAF/MEK or EGFR targeted therapies. Altogether, this argues that blocking the pathways that activate SRC kinases represents a logical strategy to reinforce the inhibition of both the RAF/MEK/ERK and EGFR/PI3K/AKT axes, and to prevent the emergence of a therapeutic resilience phenotype (**Fig. 7f**).

FDA-approved SRC inhibitors are available, however toxicity in combination with other targeted agents is a major concern. This prompted us to determine what upstream pathways activate SRC in order to potentially leverage these mechanisms as clinically actionable targets. Our data show that PGE2-signaling drives SRC activation in BRAF^V600E^ CRC. Despite decades of work on SRC, its contribution as a regulator of cancer-related inflammation has remained largely unexplored. Here, we demonstrated that BRAF^V600E^ CRC cells overcome BRAF ± MEK or EGFR therapies by upregulating a pro-survival, auto-/onco-crine, COX2 / PGE2 / GNAS / SRC / beta-Catenin signaling loop. This adaptive response was not observed in BRAF^V600E^ melanoma cells, highlighting how SRC is embedded in pre-existing signaling networks specific to BRAF^V600E^ CRC. Of note, >90% of CRC tumors also harbor alterations in the WNT signaling pathway, typically an initiating APC mutation ^40^. Kinase circuits and drug-response mechanisms are inevitably adapted to the WNT/APC-mutated background of BRAF^V600E^ CRC cells. Both the WNT and G-protein-regulated signaling networks are induced by extracellular inflammatory cues, and both share many intracellular signaling components, such as GSK3b or beta-catenin ^21,41^. This inherent predisposition of BRAF^V600E^ CRC cells to rely on inflammatory pathways to alleviate and withstand drug pressure is underscored by our finding that the SRC-relayed PGE2 signaling cascade causes therapeutic resistance (**Fig. 7f**).

The production of PGE2 is regulated by the COX2 enzyme. COX2 is a rational target, implicated in intestinal inflammation and, by association, CRC initiation and progression ^24, 42^. Although several prior clinical trials testing SRC or COX2 inhibitors in CRC patients have failed to show meaningful clinical activity ^43–48^, no trial has yet focused on BRAF^V600E^ mCRC, or evaluated these agents in combination with BRAF-targeted therapies. Recent BRAF^V600E^ mCRC clinical trials have proven the feasibility of administering three targeted therapies simultaneously ^5^, however, quadruple therapy pushes the limits of acceptability— unless the fourth therapy is inexpensive and has an exceptionally favorable side-effect profile. On both accounts, COX2 is a more attractive drug target than SRC. We showed that adding celecoxib to a current standard of care treatment, encorafenib (BRAFi) + panitumumab (EGFRi) ^5^, resulted in sustainable and significant tumor growth inhibition in BRAF^V600E^ CRC PDXs. Addition of the inexpensive, FDA-approved COX2 inhibitor, celecoxib, to BRAF-targeted therapies could be rapidly translated in patients with BRAF^V600E^ mCRC, with few anticipated sideeffects.

We attempted to determine retrospectively whether use of celecoxib or other nonsteroidal anti-inflammatory drugs as concomitant medications conferred benefit to CRC patients who participated on clinical trials testing BRAF inhibitor-based therapies, however incomplete data collection and heterogeneous dosing precluded this analysis. Additionally, while the PDX models faithfully recapitulate patients’ initial responses to targeted therapies ^6^, it is unknown whether or not they can fully predict response, especially as the mice lack a functional immune system. Furthermore, we acknowledge that there may be alternative routes to adaptive resistance, not represented by our models; in which case, it may be possible to leverage the HT-KAM platform to develop predictive biomarkers to tailor therapy most effectively.

In conclusion, our results demonstrate that SRC plays a dominant role in mediating the unresponsiveness of BRAF^V600E^ colorectal tumors to BRAF inhibitors, and suggest that SRC and EGFR act in conserved, complementary, parallel circuits that drive resistance, and can be jointly targeted to restore therapeutic sensitivity. SRC activation is non-redundant with the MAPK pathway. We determined that a COX2-inflammatory pathway drives SRC activation; the effects of inhibiting SRC were recapitulated *in vitro* and *in vivo* by targeting COX2. These results argue that drug resistance can result from a combination of pathways that are upregulated, working in concert, and interdependent on each other, such that impeding their coordinated signaling activities is necessary to overcome resistance. The HT-KAM approach can identify a finite number of key cooperative dependencies, offering a curated selection of convergent targets to evaluate. Our hope is that by expanding the scope of investigation, BRAF^V600E^ mCRC will one day become like HER2-positive breast cancer, where what was once a poor prognosis subtype with few treatment choices has been transformed into an opportunity to receive effective targeted therapy options.

## SUPPLEMENTAL FIGURE LEGENDS

**Extended Data Fig. S1.**
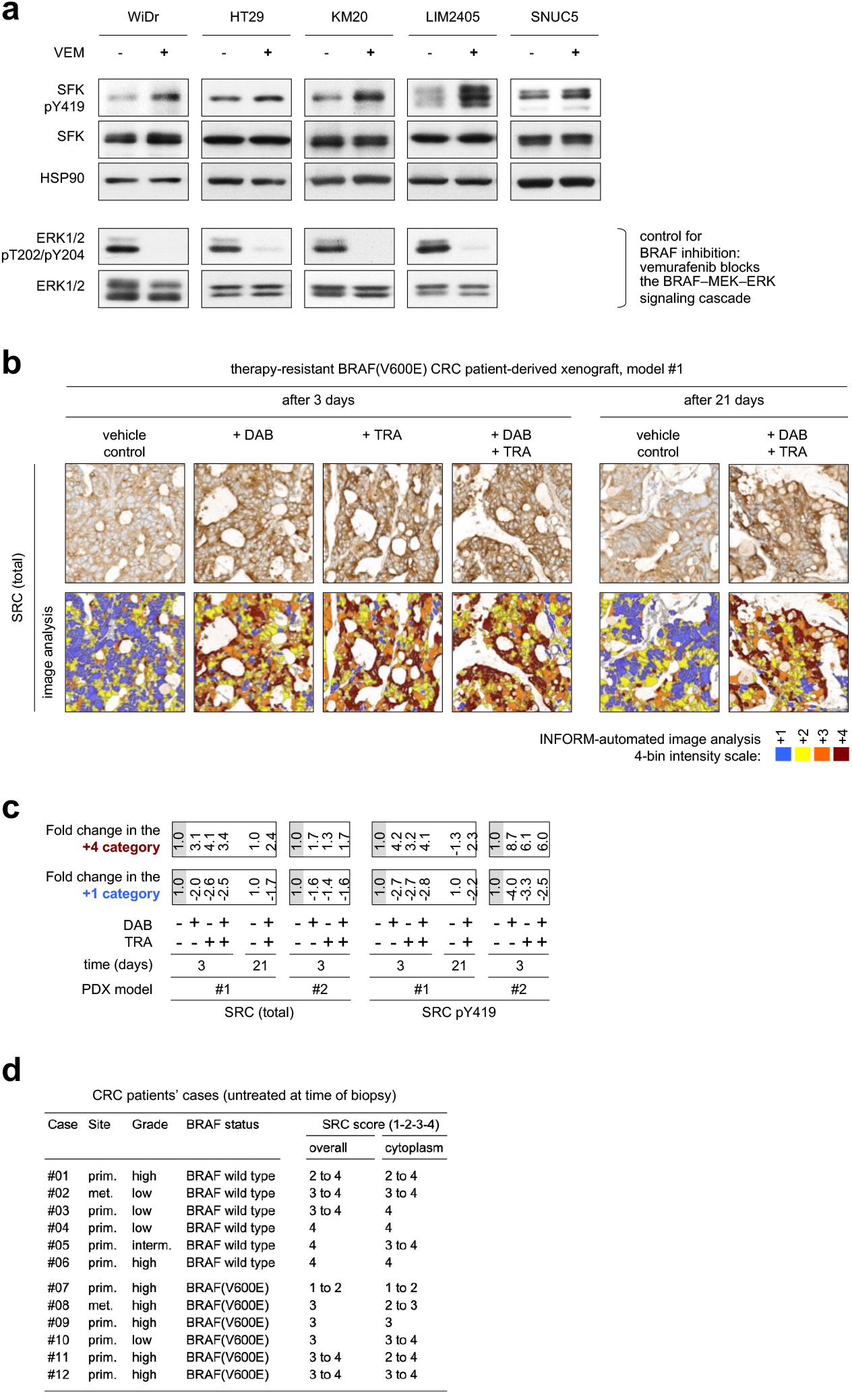
SRC is activated consequent to BRAF/MEK/EGFR inhibition in BRAF^V600E^ CRC specifically. **a,** BRAF^V600E^ CRC cell lines were treated with vemurafenib (VEM) for 7 to 8 hours. Vemurafenib was used at 1.75 uM in HT29, 2 uM in KM20, 0.15 uM in LIM2405, 2.25 uM in SNUC5, and 1.5 uM in WiDr (details of treatment conditions (concentration and time) are provided in the spreadsheet **Suppl_Table S1** and **Suppl_Table S2** in the supplementary XLS document **Source Data & Experimental Conditions**). Cell lysates were assayed by western blot with the indicated antibodies. Upper panels: SFK activation is reflected by increased phosphorylation of the SRC activation site, Y419 (pY419). HSP90 is used as loading control. Bottom panel: reduction in ERK 1/2 phosphorylation as control of BRAF inhibition. **b,** Representative IHC images showing total SRC staining intensity following treatment of a BRAF^V600E^ CRC PDX model with vehicle control, dabrafenib (DAB) and/or trametinib (TRA) for 3 or 21 days. The color-coded bottom panel highlights differences in bin intensities resulting from automated image analysis. **c,** Quantification of IHC staining intensity for total and activated SFK in two PDX models treated for 3 or 21 days with DAB ± TRA vs. vehicle control. **d,** SRC staining score by IHC in untreated patient CRC tumor specimens with or without a BRAF^V600E^ mutation, from primary (prim.) or metastatic (met.) sites.

**Extended Data Fig. S2.**
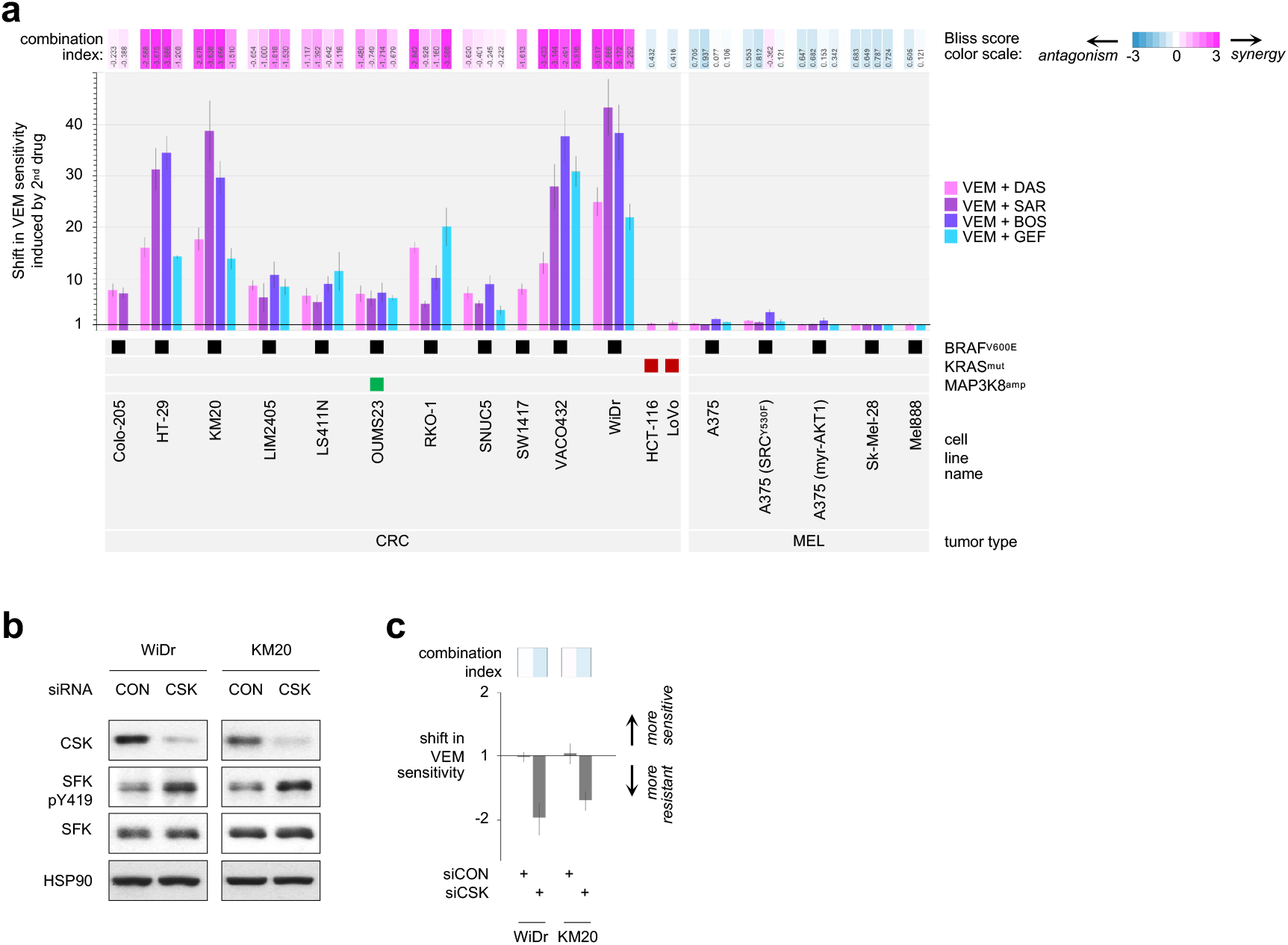
SRC kinase activity is inversely correlated with sensitivity to vemurafenib in BRAF^V600E^ CRC cell lines. **a,** Shift in vemurafenib (VEM) sensitivity, measured via cell viability assays and calculation of the combination index (CI score; top panel) upon treatment of BRAF^V600E^ or KRAS mutated or MAP3K8 amplified CRC, or melanoma (MEL) cell lines with VEM together with: a SRC inhibitor, dasatinib (DAS), saracatinib (SAR) or bosutinib (BOS), or an EGFR inhibitor, gefitinib (GEF), for three days. **b,** BRAF^V600E^ CRC cell lines were transfected with a siRNA against CSK, a negative regulator of SFKs, and siRNA Control. Cell lysates were assayed by western blot with the indicated antibodies, showing the effect of CSK depletion on SFK activation. **c,** Bar graphs representing fold-change (log scale) ± standard error for change in sensitivity to VEM with knockdown of CSK in 3-day cell viability assays. Top panel: combination index, Bliss model score; colors as in **Fig. 2b**.

**Extended Data Fig. S3.**
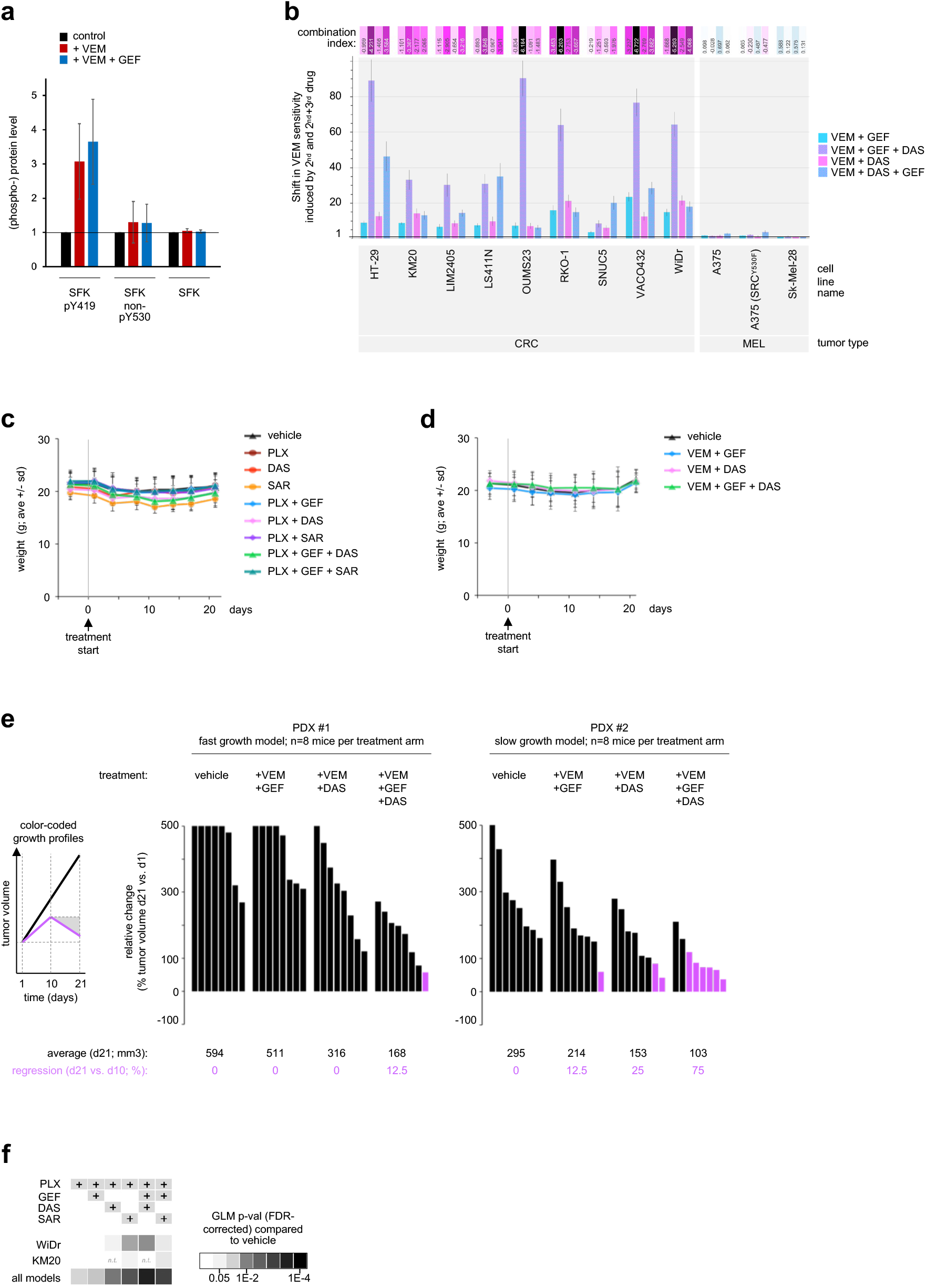
Coordinated targeting of SRC with BRAF ± EGFR improves efficacy in BRAF^V600E^ CRC cell lines and xenografts without increasing toxicity. **a,** Levels of pY419, non-pY530, and total SFK measured by western blot in BRAF^V600E^ CRC cell lines treated with vemurafenib (VEM) ± gefitinib (GEF) were quantitated. Data are normalized to control untreated per cell line and displayed as average ± standard deviation measured across HT29, KM20, LIM2405, WiDr (data provided in the spreadsheet ‘Fig 3a’ in the supplementary XLS document **Source Data & Experimental Conditions**). **b**, BRAF^V600E^ CRC cell lines were treated with exogenous PGE2. **b,** Shift in vemurafenib (VEM) sensitivity, measured via cell viability assays and calculation of the combination index (CI score; top panel) upon treatment of BRAF^V600E^ CRC or melanoma (MEL) cell lines with VEM + gefitinib (GEF) ± dasatinib (DAS) and VEM + DAS ± GEF, for three days. The addition of DAS to VEM+GEF increases sensitivity to VEM to a greater extent than the addition of GEF to VEM+DAS, highlighting the contribution of SFK and supporting that SFK activation upon VEM treatment is EGFR-independent. **c** and **d,** Mouse weight as a surrogate for toxicity following treatment of BRAF^V600E^ CRC cell line- (**c**) or patent-derived xenografts (**d**) with vehicle control or the inhibitors listed (PLX: PLX4720; SAR: saracatinib). Data is displayed as the average weight in grams ± standard deviation. **e,** Tumor growth inhibition in BRAF^V600E^ CRC PDX models following treatment with VEM ± GEF ± DAS or vehicle (control). Waterfall plots show the relative change in tumor volume: each bar represents one tumor; and the height of the bar compares the final volume at day 21 to the starting volume at day 1. Volume changes are capped at 5-fold of the starting volume (i.e. 500%). Average final tumor volumes per treatment group are indicated underneath the graph (black font). T-test p-values are indicated when p<0.05. Tumors that regressed by day 21 compared to volume at mid-treatment (i.e., day 10) are shown in purple; percentages of regressing tumors per group are indicated underneath the graph. **f,** The GLM p-values corrected for false discovery rate (FDR) corresponding to the main **Figure 3f**, are shown.

**Extended Data Fig. S4.**
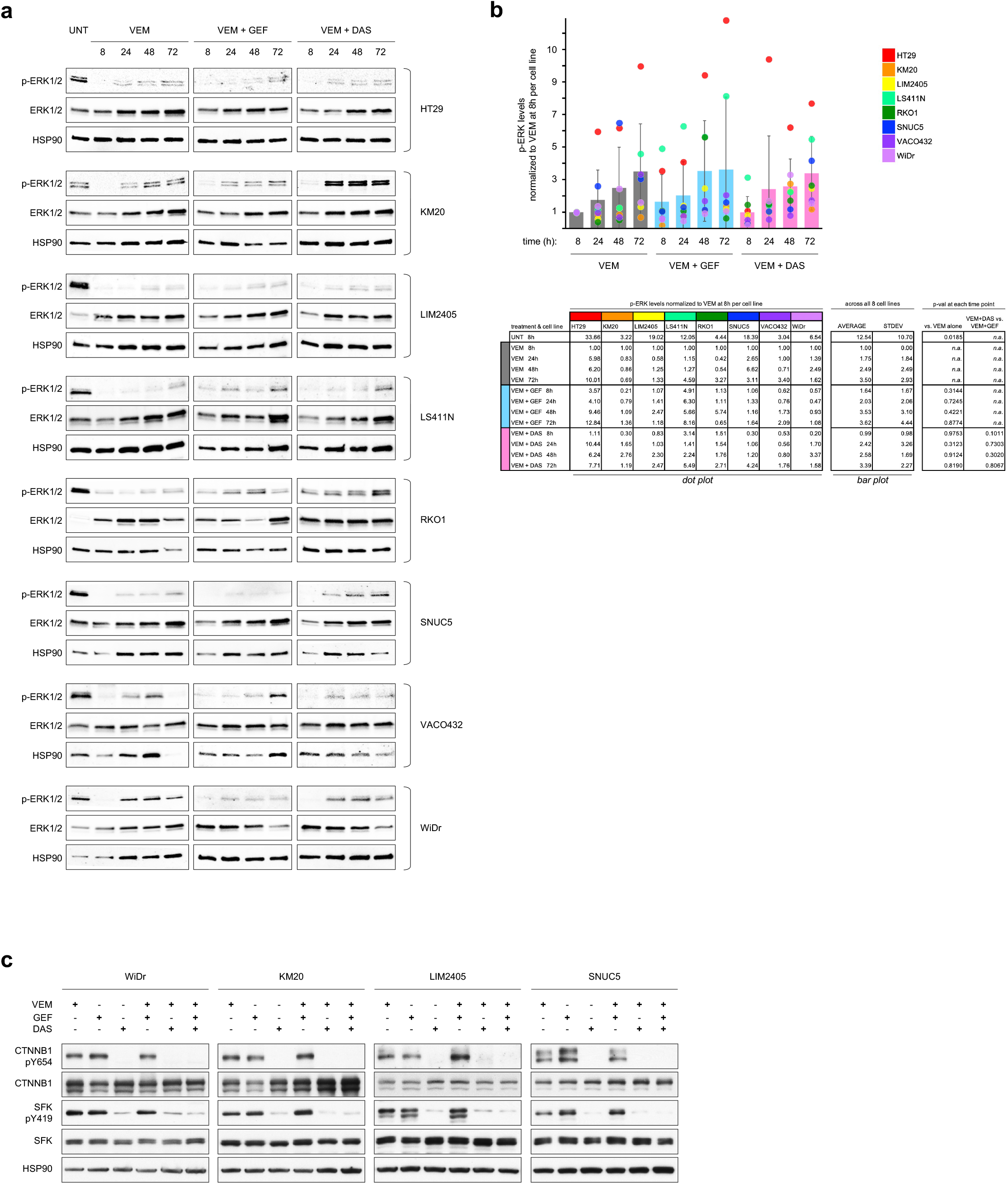
Mechanisms underlying the synergistic effects of co-targeting SRC and the BRAF ± EGFR pathways. **a,** Western blots to detect phospho-T202/Y204 ERK1/2 and total ERK1/2 in BRAF^V600E^ CRC cell lines treated with vemurafenib (VEM) ± gefitinib (GEF) or dasatinib (DAS) collected after 8h, 24h, 48h or 72h. HSP90 is used as a loading control. **b,** Quantification of western blots shown in panel (**a**). The bar plot (averages and standard deviations per treatment condition across cell lines) was overlaid with a dot plot displaying individual measurements per cell line and condition. Data are normalized to p-ERK levels after 8h treatment with VEM alone. See table below for detailed values and color codes. **c,** Western blots to detect total and phospho-Y654 beta-catenin (CTNNB1) in BRAF^V600E^ CRC cell lines treated with VEM, or GEF, or DAS, or combinations of VEM + GEF, or VEM + DAS, or VEM + GEF + DAS. The detection of phospho-Y419 and total SFK serves as a control for the effect of SFK-inhibition (with DAS).

**Extended Data Fig. S5.**
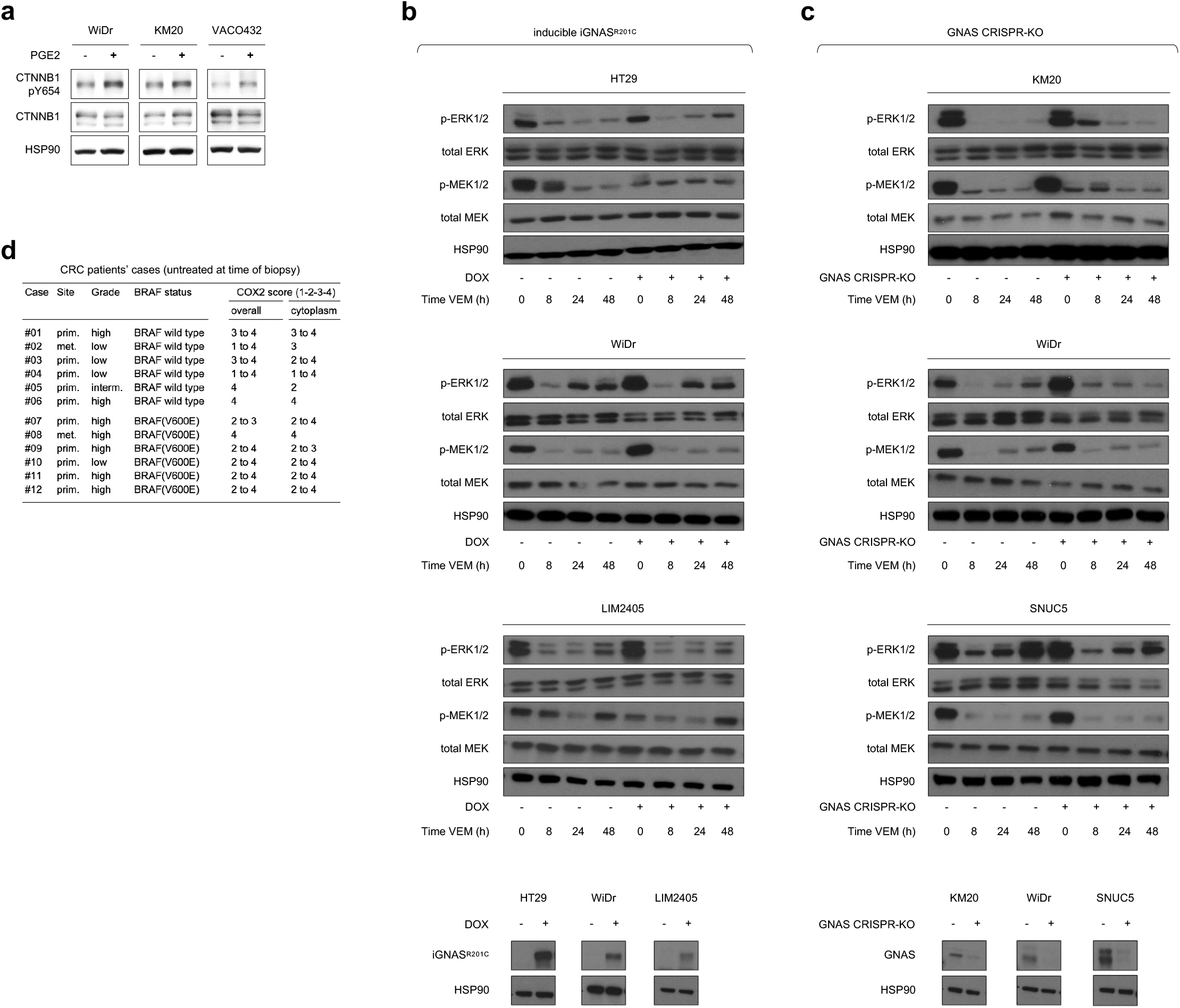
PGE2 / GNAS signaling in BRAF^V600E^ CRC cell lines and tumors. **a,** Western blots to detect phospho-Y654 of beta-catenin (CTNNB1) in BRAF^V600E^ CRC cell lines treated with exogenous PGE2. **b-c,** Western blots to detect phospho-T202/Y204 ERK1/2, total ERK1/2, phospho-S217/S221 MEK1/2, total MEK1/2 in BRAF^V600E^ CRC cell lines modified for GNAS expression, and treated with vemurafenib (VEM). **b**, BRAF^V600E^ CRC cell lines engineered with a doxycycline-inducible constitutively active GNAS construct, iGNAS^R201C^ with or without doxycycline treatment (+ or – at the bottom of each panel). **c**, BRAF^V600E^ CRC cell lines knocked out for GNAS using CRISPR (GNAS-KO or CRIPSR control; indicated as + or – at the bottom of each panel). **d,** COX2 staining score by IHC in untreated patient CRC tumor specimens with or without a BRAF^V600E^ mutation, from primary (prim.) or metastatic (met.) sites.

## Author contributions

A.R-S., C.W., B.P., C.A.D., D.B., A.P., D.P.M., D.S.S., D.J.R., V.B., E.F., Y.C.H., C.M., C.S., D.M.W. generated experimental samples, acquired data, and analyzed results. C.E.A., B.H., C.C., S.R.V. conducted pre-clinical mouse work, patient-derived xenografts, and histopathology analysis. J-P.C. conceived the kinase activity screening system and supervised the study. J.S.G., R.B., M.M.M, L.J.v.t.V., K.S., D.P.M, provided essential input for experimental designs, data interpretation, and writing. C.E.A., A.R-S., and J-P.C. wrote the manuscript. All authors reviewed the manuscript.

## Acknowledgements

We thank Eric Chow, at the Center for Advanced Technologies, UCSF, and Miki Mori for support in setting up the HT-KAM liquid assay automation, Julia Malato, Donghui Wang, Paul Phojanakong, Veronica Steri, and Debarko Banerji for their technical assistance with *in vivo* mouse studies, Jennifer Bolen and Mohammad Nasser for providing details of immunohistochemistry automation protocols, as well as the UCSF HDFCCC Histology & Biomarker Core for methods support. We thank Amgen for providing the panitumumab.

## ONLINE METHODS

### Cells lines, cell culture conditions, genetic alterations

Cell lines used in this study were purchased from ATCC or provided by Dr. R. Bernards; (i) BRAF^V600E^ CRC cells: WiDr, SNUC5, HT29, Colo-205, RKO-1, LIM2405, KM20, LS411N, VACO432, SW1417; CRC MAP3K8^amp^: OUMS23; (ii) KRAS^mut^ CRC cells: HCT116, LoVo; (iii) BRAF^V600E^ melanoma cells: A375, A375 SRC^Y530F^, A375 (myr-AKT1), Sk-Mel-28, Mel888. Cells were cultured following ATCC’s instructions or as previously described (13).

#### Transfections

siRNA transfections were performed using Lipofectamine 2000 according to the manufacturer’s instructions (Invitrogen). Cells were processed 72 hours after siRNA transfection. Control siRNAs and siRNAs targeting CSK were obtained from Dharmacon (siCON: D-001206-13-5 and D-001206-14-20; siCSK: M-003110-02).

#### Generation of knocked down SRC and control stable cell lines

For stable knockdown of SRC, the oligonucleotides containing the shRNA hairpin were annealed and then ligated into the pLKO.1 vector. The two shRNA sequences used against SRC were shSRC#1 (hairpin sequence #TRCN0000199313: 5’-GCTGACAGTTTGTGGCATCTT −3’) and shSRC#2 (hairpin sequence #TRCN0000195339: 5’-CATCCTCAGGAACCAACAATT −3’). Lentiviruses were produced by the UCSF Viracore. WiDr, KM20, LIM2405 and SNUC5 cell lines stably expressing shRNAs control and against SRC (shCON, shSRC #1, shSRC #2) were produced by transduction with the corresponding lentiviruses in the presence of 8 μg/ml polybrene (Sigma-Aldrich). After incubation in growth medium for 72 hours, cells were treated with 2μg/ml puromycin, the selection antibiotic expressed by the viral vector, to remove non-expressing cells. Puromycin-resistant cell populations were used for the various experiments described in the relevant sections.

#### Generation of GNAS lenti-CRISPR knockout cell lines

The *GNAS* lenti-CRISPR knockout cancer cell lines were generated by targeting exon 1 of the human *GNAS* locus with the CRISPR/Cas9 system as previously described (49, 50). The forward and reverse sgRNA-targeting sequences used were 5’-CACCGCTACAACATGGTCATCCGGG-3’ and 5’-AAACCCCGGATGACCATGTTGTAGC-3’, respectively. Guide sequences were provided by Aska Inoue (Tohoku University). Oligonucleotides for the sgRNAs were phosphorylated and annealed for insertion into *BsmBI* digested lenti-CRISPR v2 backbone (Addgene cat# 52961). Lentiviruses were produced by transfecting HEK293T17 cells with enveloping, packaging, and guide DNAs at a 1:2:3 ratio. Media was collected at 48 and 72 hours post-transfection. Viral particles were concentrated by ultracentrifugation at 28,000rpm for 4 hours at 4°C. Cancer cell lines were seeded on poly-lysine coated 6-well plates and transduced once the cells reached 70% confluence. Media was refreshed after 48h and cells were transduced again for 48h. Polybrene (10μg/ml) was used to enhance the transduction efficiency. Cells were selected with 1μg/ml of puromycin for 5 days.

#### Generation of Tet-GNAS active mutant cell lines

The *GNAS* active mutant was previously generated by site-directed mutagenesis of arginine 201 to cysteine (51). GNAS^R201C^ cDNA was cloned into the pENTR backbone (pDONR221, ThermoFisher Scientific, Catalog # 12536017) using the Gateway cloning BP reaction according to manufacturer protocols (Invitrogen, Catalog #11789020). Activity of the GNAS^R201C^ active mutant was confirmed by cAMP responsible element (CRE) luciferase assay (Dual-Glo Luciferase Assay System, Promega Catalog #E2920) or cAMP immunoassay (R&D Systems, Cat. #KGE002B). The GNAS^R201C^-pENTR vector was then recombined with the lentiviral vector pLVX-TetOne FLAG Puro (kindly provided by Dr. Krogan’s group, UCSF) using the Gateway LR reaction according to the manufacturer protocol (Invitrogen, Cat. #11791020). Viral particles were harvested from HEK293T17 cells and concentrated by ultracentrifugation. Cells were transduced two times for 48 hours and then selected with 1μg/ml puromycin for 5 days. Expression of GNAS^R201C^ was induced by adding 1μg/ml of doxycylcine.

### Cell extracts

For samples to be analyzed with the HT-KAM platform, cells at ~ 85% confluency were washed three times with cold PBS and lysed with freshly prepared 1X cell lysis buffer (1ml per 2.5×10^6^ cells) (10X Cell Lysis Buffer, Cell Signaling; cat.# 9803) complemented with 1X of Halt Protease & Phosphatase (100X, ThermoScientific; cat.# 1861281). Cell lysates were collected and spun down at 14,000rpm for 15min at 4°C and supernatants stored at −80°C. For samples to be analyzed by western blot, cell lysates were prepared with RIPA lysis buffer (150 mM NaCl, 0.1% SDS, 1% Nonidet P40, 1% sodium deoxycholate and 10 mM sodium phosphate, pH 7.2) supplemented with protease and phosphatase inhibitors. After clearing by centrifugation at 12,000 rpm for 10 minutes at 4°C, the lysates were analyzed as described in the respective experiments. For samples to be analyzed with HT-KAM, WiDr cells were treated as detailed in Prahallad et al, 2012 Nature (reference #13).

### Kinase inhibitors and cell treatment conditions

Inhibitors purchased from Selleck Chemicals are: bosutinib (SKI-606, cat# S1014), celecoxib (SC 58635, cat# S1261), cetuximab (cat# A2000), dabrafenib (GSK2118436, cat# S2807), dasatinib (BMS-354825, cat# S1021), gefitinib (ZD-1839, cat# S1025), PLX4720 (cat# S1152), saracatinib (AZD0530, cat# S1006), trametinib (GSK1120212, cat# S2673), valdecoxib (cat# S4049), vemurafenib (PLX4032, cat# S1267). Encorafenib was purchased from MedChem (cat# HY-15605). Panitumumab was provided by Amgen Oncology (Thousand Oaks, CA). Prostaglandin E2 (PGE2) was purchased from R&D Systems (cat# 2296).

#### Cell treatment conditions

Drug treatment conditions (concentration and time) related to western blots, ELISA, kinase activity profiles and qRT-PCR, are provided in the spreadsheet **Suppl_Table S1** and **Suppl_Table S2** in the supplementary XLS document **Source Data & Experimental Conditions**. For western blots, cells were serum starved (0.25% FBS) for 16h prior to treatment (treatment conditions provided in **Suppl_Table S2**). We placed a demarcation line where blots are not contiguous (see **Fig. 3a**, **Fig. 4a**, **Fig. 5d**, and **Fig. 5f** bottom panel).

### Cell viability assays

To assess the growth/survival response of cell lines to single or combinatorial drug treatments, we used CellTiter-Glo cell viability assay (Promega; cat# G7571). Cell culture and luminescence readouts were performed in 96- and 384-well plates after 3-day treatments. GI50 corresponds to the concentration of a given drug that causes 50% inhibition of cell growth (GI50) after 3 days of treatment. For figure panels displaying cell survival results (i.e., shift in VEM sensitivity and synergy analysis shown in **Fig. 2a,b,e, Fig. 3b, Fig. 5c,e,g, Fig. 6a,b, Extended Data Fig. S2a,c, Extended Data Fig. S3b)**, the experimental conditions for all 2-fold serial dilution treatments were started at ≥8 fold GI50 concentration, and ended at ≥0.125 fold GI50 concentration, where all concentrations were adapted/specific to each cell line and each drug. The effects of drug combinations on cell growth were assessed by calculating fold change in VEM-sensitivity and combination index (C.I.) that were experimentally measured from ≥9 individual datapoints around drugs’ GI50, i.e. at GI50, 0.5xGI50, 2xGI50 concentrations of each drug per drug combination experiment. To address the particular effects of some drug treatments on some cell lines, the choice of experimental datapoints centered around drugs’ GI50 effects was adjusted by including individual datapoints ranging from ≥GI25 to ≤GI75, leading to calculate fold change in VEM-sensitivity and combination index (C.I.) from 12 to 20 individual datapoints, as previously explained (7). For **Fig. 5c**, cells were treated with serial dilutions of PGE2 starting at 128pg/mL/12h over 3 days.

To calculate CI values in **Fig. 2a,b,e**, **Fig. 5c,e,g**, and **Fig. 6a**, we applied the Bliss Independence model (54,55), which uses experimental profiles and avoids inaccuracies that commonly occur with dose-effect curve estimation approaches. CI is calculated as CI = −log2 (Eab/(Ea*Eb)), where Ea and Eb correspond to the effects of drugs a and b alone at a given concentration, and Eab corresponds to the combined effects of drugs a and b at these same concentrations. In this model, CI>0 indicates synergistic effect, CI=0 indicates additivity effect, CI<0 indicates antagonistic effect.

To compare the effects of triple versus dual drug combinations in **Fig. 3b**, we calculated CI as follow: CI = −log2 (Eabc/(Ea*Eb)), where Ea and Eb correspond to the effects of drugs a/VEM, b/GEF alone at a given concentration, and Eabc corresponds to the combined effects of drugs a and b at these same concentrations combined with a third drug (c/DAS) at the cell line-specific GI50 of DAS. The same method was applied to analyze data shown in **Fig. 6b**.

### Colony formation assays

Colony formation assays were performed as previously described in Prahallad et al, 2012 Nature (13). In brief, to test the responses of CRC cells to different treatments, cells were plated in medium containing 10% FBS 24h prior to being washed with serum-free medium, and cultured for 24h in medium containing 0.1% serum. After low serum incubation, cells were treated with drugs for 30 min and stimulated by 10% FBS.

### Kinase activity mapping assay

This high throughput kinase-activity mapping (HT-KAM) platform uses arrays of peptides that act as sensors of phosphorylation activity (7). The phospho-catalytic signature of samples is established from simultaneously occurring ATP-consumption tests measured in the presence of individual peptides that are experimentally isolated from each other. Assays are run in 384 well-plates, where each experimental well contains one peptide. The final 8μL reaction mixtures per well contain: (i) kinase assay buffer (1X KaB: 2.5mM Tris-HCl (pH7.5), 1mM MgCl_2_, 0.01mM Na_3_V0_4_, 0.5mM β-glycerophosphate, 0.2mM dithiothreitol (DTT), prepared daily; 10X KaB Cell Signaling cat.# 9802), (ii) 250nM ATP (prepared daily with 1X KaB; Cell Signaling cat.# 9804), (iii) 200μg/ml 11-mer peptide (lyophilized stocks originally prepared as 1mg/ml in 1X KaB, 5% DMSO), and (iv) samples made from cell at ~10μg/ml total protein extract. Samples are kept on ice and diluted in 1X KaB <30min before being used. Controls with no-ATP, or no-peptide, or no-sample as well as ATP standards are run side-by-side within each 384-well plate. High-throughput liquid dispensing of all reagents is performed using a Biomek^®^ FX Laboratory Automation Workstation from Beckman Coulter. All reagents are kept on ice and plates on cold blocks until enzymatic reactions are started. Once the dispensing of the reaction mixtures is complete, the plates are incubated for 1h at 30°C. ATP is detected using Kinase-Glo revealing reagent (Promega; cat.# V3772), which stops the activity of the kinases and produces a luminescent signal that directly correlates with the amount of remaining ATP in the samples. Luminescence is acquired using the Synergy 2 Multi-Mode Microplate Reader from BioTek. Luminescence data are inversely correlated with the amount of kinase activity. For a more detailed description of the peptide sensors design, sequence and connectivity between peptides and kinases, as well as data normalization steps and analysis, refer to: (7, 9, 10, 52, 53). The activity of kinase enzymes is derived from their respective subset of biological peptide targets included in the assay.

### Antibodies and western blotting

For western-blot, samples were denatured by boiling in 1X Laemmli buffer and run on an 8% SDS-PAGE gel. After transfer onto a PVDF membrane and blocking with 3% BSA, the membranes were incubated with primary antibodies overnight at 4°C, washed 3 times with TBST (Tris-buffered saline containing 0.05% Tween-20), incubated with secondary antibodies for 1 hour at room temperature and developed using chemoluminescence (cat# 32209 from Pierce).

Antibodies anti-HSP90 (H-114, cat# 7947) and anti-phospho-beta-catenin (CTNNB1 Y654, cat# 57533) were procured from Santa Cruz Biotechnologies. Anti-phospho Src (Y419) (cat#AF2685) used for IHC was obtained from R&D Systems. Anti-phospho Src Family (Y416) (D49G4, cat# 6943) used for western-blots, anti-non-phospho-Src (Y527) (cat# 2107); anti-Src (32G6, cat# 2123) used for western-blots and anti-Src (cat#2109) used for IHC, anti-phospho-p44/42 MAPK (ERK1/2 T202/204, cat# 9101); anti-p44/42 MAPK (ERK1/2, cat# 9102), anti-phospho-MAP2K1/2 (MEK1/2 T202/204, cat# 9121); anti-MAP2K1/2 (MEK1/2, cat# 9122) and anti-CSK (C74C1, cat# 4980) were from Cell Signaling. Anti-Gsalpha-Subunit (GNAS, cat# 371732) was obtained from CalBiochem (now Millipore/Sigma), anti-COX2 was purchased from Spring Bioscience (cat# M3210), anti-Gs alpha subunit, C-terminal (385-394) antibody was purchased from Calbiotech (cat# 371732), anti-beta-catenin was purchased from BD Transduction Laboratories (CTNNB1, cat# 610153). Secondary antibodies Horseradish peroxidase (HRP)-conjugated were from GE Healthcare (rabbit cat# LNA934).

In an inactive form, SRC is phosphorylated at tyrosine Y530 (Tyr530 in mammalian c-Src; Tyr527 in chicken c-Src) near the C-terminus of SRC. Phosphorylation at Y419 (Tyr416 in chicken c-Src) is associated with SRC activation. The SFK family encompasses 11 members in humans by the Manning classification (Manning et al., 2002) and the numbering for those two key tyrosines varies for some of the other family members. SRC antibodies used in western-blots cross-react to certain extent with other members of the SFK (data not shown).

### ELISA assay

To detect secreted PGE2, conditioned media of BRAF^V600E^ CRC cell lines treated with vemurafenib (VEM) ± gefitinib (GEF) was collected. Particulates were removed by centrifugation and samples were processed according to the manufacturer protocol (R&D, Catalog KGE004B).

### RNA extraction, cDNA synthesis and quantitative real-time PCR (qRT-PCR)

RNA was purified using TRIzol followed by DNAse RNase-free digestion (Qiagen). cDNA synthesis was performed using the High Capacity cDNA Reverse Transcription kit (ABI). Primer efficiencies were assessed by serial dilutions. qRT-PCR reactions were performed in QuantStudio™ 5 Real-Time PCR System using SYBR-Green Power master mix (ABI) with default cycling conditions; results were analyzed with the QuantStudio™ 5 analysis software. All mRNA levels were assayed in quadruplicates; dissociation curves were checked and products were run in agarose gels to confirm amplification of only one product. Relative mRNA levels of beta-catenin target genes (*MYC, AXIN2, ASCL2, S100A6, LEF1, NOTCH2, SP5*) were calculated by the 2^^(-ΔΔCt)^ method using *ACTB* and *UBC* as controls. The sequences (5′➔3′) of primers we used to measure the mRNA levels of beta-catenin target and housekeeping genes are provided in the spreadsheet **Suppl_Table S3**. For statistical analysis of qRT-PCR results, we used a t-test, two-sample equal variance (with a two-tailed distribution), to determine the significance of differences in gene expression.

### Automated immunohistochemistry procedure

To identify changes in protein expression in tumors from PDXs treated or not with dabrafenib and/or trametinib (**Fig. 1e, Fig. 4h, Extended Data Fig. 1b**), and tumors from patients (**Extended Data Fig. 1d, 4a**), immunohistochemistry staining was preformed using the Ventana DISCOVERY ULTRA autostainer system hosted at the UCSF Histology and Biomarker Core. A critical advantage of the fully automated DISCOVERY ULTRA pipeline is that, once an IHC protocol is set, all parameters and workflow are automated and repeated identically, thus allowing for all biospecimens to be processed in the exact same way, which further allows for automated image processing and comparative analysis.

Ventana reagents where used, except where noted, according to manufacturer’s instructions. Specific settings for each antibody were programed on the DISCOVERY ULTRA, as detailed below. Tissues known to express the marker of interest were included as positive control with each staining run, and during optimization the same tissue was used following the optimized protocol with primary antibody omitted to generate a background reference negative control slide.

Briefly, slides where sectioned at 4 μm thickness, mounted on positively charged slides, and air dried. To increase tissue adhesion slides were baked in oven at 60°C for a minimum of one hour to a maximum of 24h. De-paraffinization was done on the DISCOVERY ULTRA in three cycles of 8min each in EZ Prep solution warmed to 72°C. Antigen retrieval was performed at high temperature, between 95°C and 100°C, in Cell Conditioning 1 Solution (similar to EDTA) for 4 to 92min as required by the tissue and antibody combination. Before primary antibody application, inhibitor (specifically Inhibitor CM from the DAB kit) was applied and incubated for between 8 and 20min. If additional blocking was required to reduce background this was followed by DISCOVERY Goat Ig Block RUO product number 760-600 for 4 to 16min. Primary antibody was diluted in Discovery Ab Diluent. Species-specific secondary antibody, either OmniMap or HQ and enzyme conjugate, was applied and incubated with low heat between 36 and 37°C. DAB from the DISCOVERY ChromoMap DAB RUO kit was selected as the chromogenic detection for which the Discovery Ultra hard codes the incubation settings. Hematoxylin nuclear counterstain (Cat# 760-2021) was applied for 4min followed by Bluing reagent for additional 4min. Slides were washed and dehydrated according to Ventana standard operating procedure and coverslipped using 0.17mm thick glass coverslips and Cytoseal XYL mounting media Richard-Allan Scientific, Cat.# 22050262.

To detect phospho-Y419 SRC, phospho-Src (Y419) EGFR rabbit polyclonal supplied by R&D Systems (Cat# AF2685) was used at a titration of 1:50. Cell conditioning in CC1 was performed at 95°C for 64min. Inhibitor was applied for 16min. The primary antibody was incubated at 37°C for 32min. Anti-Rabbit HQ was used as the secondary antibody and incubated at 37°C for 16min followed by the anti-HQ-HRP enzyme conjugate which was applied for 16min. DAB was used as the chromogenic detection with hematoxylin selected for the counterstain (as detailed above).

To detect total SRC, TOTAL SRC (36D1) rabbit monoclonal manufactured by Cell Signaling Technology (Cat# 2109) was used at a titration of 1:800. Cell conditioning in CC1 was performed at 95°C for 32min. Inhibitor was applied for 16minutes. The primary antibody was incubated at 37°C for 32min. OmniMAP anti-rabbit was used as the secondary antibody and incubated at 37°C for 12min. DAB was used as the chromogenic detection with hematoxylin selected for the counterstain (as detailed above).

To detect COX2, COX2 (SP21) rabbit monoclonal antibody manufactured by Abcam (Cat# ab16708) was used at a titration of 1:100. Cell conditioning in CC1 was performed at 95°C for 64min. Inhibitor was applied for 16min. The primary antibody was incubated at 37°C for 32min. OmniMAP anti-rabbit was used as the secondary antibody and incubated at 37°C for 8min. DAB was used as the chromogenic detection with hematoxylin selected for the counterstain (as detailed above).

### Automated processing and analysis of IHC images

Immunohistochemistry images were processed with the inForm Tissue Finder software (AKOYA Biosciences). The feature recognition algorithms available in inForm, automate the detection and segmentation of tissues and cells, as well as the quantification of immuno-staining intensities (**Fig. 1f, 5i, Extended Data Fig. S1c**). Automation provides consistent, reproducible results and enables comparative studies across specimens / samples / images.

The image processing workflow is a machine-learning process defined by, and adjusted through, an iterative sequence of user-trained modules and variables. The spectral components of each image are first unmixed using a spectral library. Next, all cells composing the immuno-stained tissue image are detected and annotated using tissue- and cell-segmentation modules. These steps locate individual cellular objects by identifying the nuclei from the spectrum corresponding to the hematoxylin stain. Based on nucleus boundaries and definable cytoplasmic/cell membrane features, the inForm software finds and draws the boundaries of each cell. This enables systematic quantification of immuno-staining intensity for each individual cell composing the tissue and across the different tissue types (e.g. tumor vs. stroma). The pattern-recognition detection/segmentation and the immuno-staining intensity scoring of tissues/cells is initially run on 3-to-5 images, and then reiterated using an additional set of 15-to-25 randomly chosen images to further finetune all parameters and validate consistency of processing across images and visualization output. Once the ‘training’ of the inForm algorithm is considered final, batch processing of all images is applied, allowing for systematic processing with identical parameters across 100’s of images, and generating results fully comparable between all individual cells from all tissues and biospecimens.

To quantify protein expression at the single cell-level in tumor areas, we automated the scoring of immuno-staining intensities using a binning tool available in the inForm software. The same binning thresholds were used for each antibody (e.g. same 4-bin intensity levels across treatments and across PDXs for SRC (total) to provide comparable results between conditions and tumor cases). However, the thresholds were specifically adapted to each antibody (i.e. different 4-bin intensity levels for SRC vs. SRC pY419 vs. COX2).

It may be noted that the IHC-intensity binning process tends to underestimate both the lowest (0/blue) and highest (+3/brown) intensities per cell. As well, the tissue segmentation tends to miss tissue areas where cells displaying very low immuno-staining intensity (which is inherent to the rules of the algorithms used to train the software to automatically differentiate between tissue types; e.g. tumor, stroma, empty).

### Patient-Derived Xenograft (PDX) models and BRAF^V600E^ CRC cell line xenograft studies

PDX models were established from consenting UCSF patients’ tumor biopsy samples as previously described (4, 11). All patients went on to receive dabrafenib + trametinib as part of a clinical trial (4). JAX NOD scid gamma mice bearing subcutaneous PDXs were randomized into vehicle or treatment groups when tumor volumes reached 100-150 mm^3^ with rolling enrollment. Mice were treated with vehicle (0.1% Tween-20 or 0.5% hydroxypropyl methylcellulose and 0.2% Tween-80) or targeted therapies for 21 days (**Fig. 3d-e**, **Fig. 6d**) or up to 60 days (**Fig. 7a-b**) in the University of California San Francisco Preclinical Therapeutics Core (San Francisco, CA).

Inhibitors administered by oral gavage (PO) were purchased from Selleck Chemicals (Houston, TX) and dosed daily (QD) as follows: celecoxib 50 mg/kg PO, QD; encorafenib 20 mg/kg PO, QD (see Krepler et al., 2016 Clin.Can.Res. (61)); dabrafenib 30 mg/kg PO, QD; dasatinib 20 mg/kg PO, QD; gefitinib 50 mg/kg PO, QD; saracatinib 25 mg/kg PO, QD; trametinib 0.6 mg/kg PO, QD; vemurafenib 50 mg/kg PO, QD. Panitumumab (used in **Fig. 7a-b**) was provided by Amgen Oncology (Thousand Oaks, CA) and was used at 200 μg by intraperitoneal injection twice weekly. For WiDr and KM20 cell line xenograft studies (see **Fig. 3d**), respectively 8 and 7 mice per treatment group were dosed daily (QD) with a combination of the following inhibitors: PLX4702 (50mg/kg/day orally), gefitinib (50mg/kg/day orally), gasatinib (50mg/kg/day orally), saracatinib (25mg/kg/day orally).

Mice were monitored for signs of toxicity (e.g. weight loss) and tumor size was evaluated twice per week by digital caliper measurements. The 15% body weight reduction threshold for holding drug was not met. The same procedures were followed for cell-line derived xenograft models. All protocols were approved by the Institutional Animal Care and Use Committee (IACUC).

To test the significance of the changes in tumor volume over time between treatment arms and vehicle in **Fig. 3d-i**, **Fig. 6d,f,g**, and **Fig. 7a-b**, we use the following statistical tests: Student t-test, generalized Linear Model (GLM), GLM p-values corrected for false discovery rate (FDR). Besides plotting the relative tumor volume in **Fig. 3d-e**, **Fig. 6d**, and **Fig. 7a-b**, we also calculated effect size measured as the GLM standard coefficient. GLM was applied to each tumor model separately, or combined. In **Fig. 7b**, none of the mice treated with ENC alone were available at the last two time points (euthanized due to tumor size), which is why no p-value could be calculated and is represented as an ‘x’ in the table underneath the plot. All raw and relative tumor volumes (and number of mice per group) can be found in the spreadsheets Fig 3d, Fig 3e, Fig 6d, Fig 7a, Fig 7b of the **Source Data & Experimental Conditions** document.

### Patient specimens

Tumor specimens were used in research following patient consent and approval by the UCSF Institutional Review Board (# 12-09139 and 13-2574).

### Statistics and Reproducibility

We provide general information on how statistical analyses of data were conducted, and general information on reproducibility of experiments.

For the analysis leading to the heatmap in **Fig. 1a**, the average value of ATP consumption in sample-containing wells measured across 228 peptides and 14 peptide-free controls was used for internal normalization for each experimental run (i.e. mean-centering value established from 242 datapoints/wells per 384-well plate; previous explained in references (7, 9, 10)). Other normalization schemes were used for further analysis and cross-validation (i.e. (i) subset of 14 peptide-free control wells (i.e. cell extract alone), or (ii) subset of 16 Y/S/T-free peptides, or (iii) subset of 63 reference peptides). The activity per-peptide was then calculated as the difference in ATP consumption between each peptide and the internal mean. ATP consumption measurements associated with each peptide were then averaged across all biological and technical replicates. Next, phosphorylation activity profiles across all individual 228 peptides were compared between treated (VEM or VEM+GEF or VEM+CET) and control (UNT) (calculated as the difference in ATP-consumption). Finally, phospho-catalytic activity signatures measured across the 228-peptide sensors were subjected to unsupervised hierarchical clustering. Phospho-catalytic activities are color-coded based on the relative level of activity measured in presence of each peptide for each treatment, from blue for ATP consumption lower than then one measured in untreated cells, to white ATP consumption for equal to the one measured in untreated cells, to red for ATP consumption higher than then one measured in untreated cells.

For the analysis to generate heatmaps in **Fig. 1b** (and related plots in **Fig. 1c-d**), the activity of kinases was calculated as the average of the phosphorylation activities measured in presence of their respective biological peptide subsets (i.e., derived from values/calculations used to generate **Fig. 1a**). Here, we systematically converted peptide-phosphorylation profiles into kinase activity signatures for individual kinases and kinase families that were detected with ≥3 distinct biological peptide sensors and available in the peptide library.

In **Fig. 3d-e** and **Fig. 7c-d**, changes in tumor volume at each time point are shown as the mean volume of all tumors per treatment arm (+/- standard error (SE)). Student t-test comparing treatment versus vehicle, or between drug treatment arms, are shown.

In **Fig. 6d**, changes in tumor volume converted to cumulative tumor volume, are shown for all individual tumors per treatment arm at day-21 (final time point to assess and compare the efficacy of different drug combinations). Significance comparing treatment versus vehicle at day-21 were calculated, and p-values (Student t-test) are shown in the plot.

In **Fig. 3f-i**, **Fig. 6f-g** and **Fig. 7c-d**, we used a generalized linear model (GLM) to analyze profiles of tumor growth. We first zero-normalized the data compared to control vehicle per model and per time point. We then ran a GLM (available in R) on the normalized data to test association of change in tumor volume between treatment and vehicle across all time points. The effect size of each treatment compared to vehicle was calculated (i.e., GLM standard coefficient) along with significance (i.e., False Discovery Rate (FDR)-corrected GLM p-values). This GLM approach and data normalization also allowed us to compare the effects of different treatments (e.g. with or without dasatinib (**Fig. 3h-i**); with or without celecoxib (**Fig. 6g, 7d**)), as well as to combine the distinct models (e.g., all cell line tumor xenografts (**Fig. 3f,h**), or all PDXs (**Fig. 3g,i**, **Fig. 6f-g** and **Fig. 7cd**)) to measure the overall effect of drug combinations and their significance.

Other statistical and predictive methods to compare sample groups and reproducibility between signatures include unsupervised or semi-supervised hierarchical clustering using Euclidean distance or (Absolute) Correlation (centered or uncentered) and Ward linkage or complete or average linkage to group phospho-activity signatures based on their similarities or differences; False Discovery Rate (FDR/BH)-corrected or not Student t-test, FDR-corrected or not Wilcoxon rank sum test (p-val<0.05); generalized linear model (GLM).

## Data availability

Data supporting the findings of this study are available from the corresponding author on reasonable request.

